# The wheat VIH2-3B, a functional PPIP5K controls the localization of fasciclin-like arabinogalactan protein

**DOI:** 10.1101/2024.09.24.614694

**Authors:** Anuj Shukla, Reshma Gopal, Riya Ghosh, Ankur Chaudhuri, Kanupriya Agrwal, Rahul Tanwar, Henning Jacob Jessen, Debabrata Laha, Ajay K Pandey

**Author notes:** Corresponding author: Dr. Ajay K Pandey, Scientist-F, **Email:**.

## Abstract

Inositol pyrophosphates (PP-InsPs) are important signalling molecules that participate in multiple physiological processes across a wide range of eukaryotes. Metabolic pathway kinases (VIP1/VIHs) leading to the production of PP-InsPs are now well characterized in yeast and plants. Previously, the wheat (*Triticum aestivum* L.) inositol pyrophosphate kinase (TaVIH2) was shown to encode a catalytic active kinase domain. Heterologous expression of TaVIH2 in *Arabidopsis thaliana* was shown to enhance drought tolerance by modulating the cell composition. In this study, we attempted to identify the interacting protein targets of wheat VIH2-3B using a yeast two-hybrid (Y2H) cDNA library screen, which led to the identification of 52 putative interactors that are primarily involved in cell wall-related functions. Notably, fasciclin-like arabinogalactan protein (FLA7), a glycosylphosphatidyl inositol (GPI)-anchored protein, emerged as the most frequently interacting partner. Further analysis using pulldown assays validated the interaction between TaVIH2-3B and TaFLA7 in vivo. Using the reporter fusion studies, we observed the localization of TaFLA7 to be a plasma membrane and this localization of the TaFLA7 was perturbed in the yeast vip1Δ strain. The expression of TaVIH2-3B bearing PPIP5K enzymatic activity in yeast mutants rescued the level of IP_8_ and restore the localisation of the TaFLA7 to the membrane. Expression analysis of TaFLA7 revealed a differential expression response to drought in wheat shoot tissues. TaFLA7 was also found to be highly expressed during grain development, particularly in the endosperm and seed coat during grain maturation. Taken together, these findings highlight the potential role of TaVIH2 in cell wall remodelling and stress response pathways, offering new insights into the functional roles of VIH proteins in plants.

## 1. Introduction

Inositol pyrophosphates (PP-InsPs) are high-energy molecules that are ubiquitous in various eukaryotic organisms and serve as critical cellular signaling entities (Szijgyarto et al. 2011). Over the last two decades, inositol pyrophosphate signalling has gained significant attention and has been proven to regulate different cellular functions such as regulating telomere length, DNA repair, phosphate homeostasis, vesicular trafficking and protein phosphorylation (Saiardi 2004; Fridy et al. 2007; Lee et al. 2007; Azevedo et al. 2009; Thota and Bhandari 2015; Shears 2015; Sanchez et al. 2019; Pipercevic et al. 2023; Chabert et al. 2023). Researchers have identified two distinct classes of inositol pyrophosphate synthases, inositol hexakisphosphate kinases (referred to as IP6Ks or Kcs1-like proteins) and diphosphoinositol pentakisphosphate kinases (known as PPIP5Ks or Vip proteins) (Choi et al., 2007). The latter kinases are primarily classified based on their catalytic specificity towards various positions on the inositol ring with a ATP-grasp domain at the C-terminal (Mulugu et al. 2007).

Genes encoding VIH-type PP-InsP synthase could be identified in green algae and embryophytes (Desai et al. 2014; Laha et al. 2015). *Arabidopsis* VIH proteins catalyze the synthesis of the inositol pyrophosphate, InsP_8_ and VIH-derived InsP_8_ has been linked with diverse physiological processes such as plant defence and phosphate homeostasis (Laha et al. 2016; Jung et al. 2018; Dong et al. 2019; Zhu et al. 2019; Ried et al. 2021; Riemer et al. 2021; Land et al. 2021). It was further shown that InsP_8_ binds to SPX1 protein and controls its interaction with the phosphate starvation response 1 (PHR1), a central regulator of phosphate starvation (Jung et al. 2018; Dong et al. 2019; Ried et al. 2021). Based on this evidence, one could speculate that PP-InsPs are associated with different physiological stress response signalling cascades.

Although the level of PP-InsPs seems to be important in asserting such function, it remains to be studied if the responsible kinases, especially plant VIH could interact physically with the components of the specific signalling system. One consequence could be local enrichment of PP-InsP species in the vicinity of their target proteins. There are multiple reports that suggests the employment of such gating of PP-InsP to its target proteins through physical interaction of the responsible PP-InsP synthase (Rao et al. 2014; Zhou et al. 2021; Laha et al. 2022; Bhandari et al. 2024). A comprehensive interaction study for the HIV protein is lacking in plants, which could provide new insights into how its signalling output is regulated. Multiple VIH proteins were identified in hexaploid wheat ((*Triticum aestivum* L.) with diphosphoinositol pentakisphosphate kinases (PPIP5K) like enzymatic activity (Shukla et al. 2021). Notably, when wheat VIH homolog (TaVIH2-3B) was expressed heterologously in *Arabidopsis*, it conferred tolerance to drought conditions (Shukla et al. 2021). Further, it was shown that VIH2 overexpressed lines show reprogramming of genes involved in cell-wall-related metabolic pathways. At the metabolome level, these transgenic lines showed high accumulation of cell-wall cementing polysaccharides such as arabinoxylan and arabinogalactan (Shukla et al. 2021). Although the role of VIH was established in imparting drought tolerance in plants, it remains to be dissected if VIH could be involved at the protein level.

Using a yeast two-hybrid (Y2H) screening system, we identified multiple interactors of TaVIH2-3B. Further, a detailed characterisation was done for one of the interactors referred to as FLA7, which is involved in the cell wall maintenance process. Using biochemical experiments and a yeast genetic screen, we demonstrated that the presence of VIH-type PP-InsP synthase is required for FLA7 localisation. This study expands our knowledge of the functioning of the plant VIHs towards new roles, including cell wall homeostasis.

## 2. Materials and methods

### 2.1. Wheat cDNA library construction and yeast two-hybrid screening

Total RNA extracted from the wheat seedlings was used to purify mRNA. mRNA was isolated by a plant mRNA isolation kit (Sigma). The mRNA concentration and purity were checked by nanodrop and 0.8% (w/v) agarose gel, then visualized by ethidium bromide staining under UV transilluminator. Multiple batches of mRNA were pooled together to prepare the cDNA library, and single-strand cDNA was constructed as described in the materials and methods section. Next, the double-strand cDNA was prepared through long-distance PCR, and aliquots were analyzed on the 1.2% agarose ethidium bromide gel. Finally, the ds-cDNA was first purified using CHROMA SPIN+TE-400 columns. For yeast-two-hybrid experiments (Y2H), the wheat cDNA library was prepared by the co-transformation of the pGADT7-Rec vector (containing SV40 Nuclear Localization Sequence) provided in the kit (Matchmaker Gold Yeast Two-Hybrid System, Takara Bio, USA) and purified cDNA prepared in the Y187 yeast strain. The transformation mixture was diluted (10^-1^ and 10^-2^), and 100 μl of each dilution was plated on the SD-L agar plates to select independent clones, indicating the readiness of our library’s complexity. In total, approximately 1.2 million independent transformants were screened, corresponding to about 5× the estimated complexity of the wheat cDNA library, ensuring a saturated screening for comprehensive identification of potential interacting partners. The full length TaVIH2 cDNA was cloned in the pGBKT7 vector using Sma1 and Not1 restriction sites.

### 2.2. Quantitative real-time PCR

To quantify the gene expression in roots, qRT-PCR was performed using the QuantiTect SYBR Green RT-PCR Kit (Qiagen, Germany). The gene-specific primers capable of amplifying the genes mentioned in TableS1 were carefully designed using Oligocalc software. The cDNA dilutions to be used were optimized by running a semi-quantitative PCR for 22-25 cycles using Thermo Scientific™ DreamTaq™ Green PCR Master Mix (2X). The qRT-PCR reaction was set up in a 96-well plate by mixing the reaction components including 1X QuantiTect SYBR Green RT-PCR Master Mix, appropriate cDNA dilution, 900nM of forward and reverse primers and MQ to final volume of 10μl in each well and the plate was run in 7500 Fast Real-Time PCR System. 4-5 technical replicates for each set of primers and a minimum of 2 biological replicates were used to validate the experiment. ADP-ribosylation factor gene (TaARF) was used as an internal control in all the expression studies (Meena et al. 2024). The Ct values obtained after the run were normalized against the internal control and relative expression was quantified using 2−ΔΔCT method (Livak and Schmittgen 2001).

### 2.3. Bioinformatics and molecular docking analysis

Intrinsic disorder region (IDR) percentage, length and position of human (*Homo sapiens*) and plant-encoded PPIP5K-like proteins were done using ESpritz (ver1.3) (Walsh et al. 2012). In silico protein-protein interaction was anlayzed using STRING analysis (Szklarczyk et al. 2023). For gene expression studies the expression browser from https://www.wheat-expression.com was used (Ramírez-González et al. 2018). Wheat eFP Browser (Triticum aestivum) cv. Azhurnaya developmental time course was used for the analysis of the gene expression at different Zadoks scale (Z) levels, which were summarised as TPM from the transcript to the gene level using tximport v1.2.0. Three biological replicates, consisting of five individual plants each. were sampled at the developmental times and tissues mentioned.

The complete protein sequences of FLA7 homoeologs FLA7-2A (TraesCS2A02G165600) and FLA7-2D (TraesCS2D02G172900) were obtained from the EnsemblPlants (https://plants.ensembl.org/Triticum_aestivum/Info/Index), followed by the removal of their transmembrane regions (TMD) to generate a truncated form of the proteins. AlphaFold2, accessed via the Neurosnap Online Bioinformatics server (https://neurosnap.ai/overview), was utilized for computational 3D structure prediction of native and truncated proteins (Yang et al. 2023). The three-dimensional (3D) structures of myo-inositol hexakisphosphate (InsP[), diphospho-myo-inositol pentakisphosphate (5-InsP[), and bis-diphospho-myo-inositol tetrakisphosphate (1,5-InsP[) were procured from the PubChem database (https://pubchem.ncbi.nlm.nih.gov/) in .SDF format. Molecular docking analyses were executed using CB-Dock2 (https://cadd.labshare.cn/cb-dock2/index.php), identifying favourable binding sites and key interacting residues (Yang et al. 2022). The protein-ligand complexes were further examined using PyMOL v3.0.0, delineating secondary structure elements and polar residues within the binding interface. Protein-ligand interactions were graphically rendered in Adobe Illustrator, accentuating critical residues and summarizing docking insights.

### 2.4. Co-immunoprecipitation and Western analysis of interacting proteins

For co-immunoprecipitation assays, respective yeast cultures were grown in SD medium with appropriate amino acids dropout. Each independent primary yeast culture [containing construct for cMYC-TaVIH2, HA-FLA7, cMYCTaVIH2+HA-FLA7 and Y187 (Wild type), respectively] was grown overnight at 30 °C upto an OD_600_ of 1. The secondary culture was incubated for 6 hours, cells were washed and the lysate was prepared using glass beads in buffer (1% SDS, 100 mM NaCl,100 mM Tris-Cl, 1 mM EDTA, 2% Triton and 1mM protease inhibitor (100X Halt protease inhibitor, Thermofisher, U.S.A.). Equal amounts of each lysate was incubated overnight at 4 °C with 100% of Protein-G agarose beads and 2 μl of anti-c-MYC antibody. The beads were subsequently washed three times with lysis buffer and centrifuged for 30 sec at 800 x g. 50 μl of loading dye was added to the washed beads and heated at 65 °C for 10 min. The elutant was resolved through SDS-PAGE and transferred to the PVDF membrane. Blot was then separated into two parts to detect TaVIH2 and TaFLA7 separately using the respective antibodies. Respective primary antibodies were used for probing (mouse Anti-c-MYC and rabbit anti-HA with 1:2000 dilution). After washing the blots with TBST, they were treated with the secondary antibody (Goat Anti-Mouse IgG (H + L); and Goat Anti-Rabbit IgG (H + L) with 1:5000 dilution. After subsequent washing with TBST, the blot was developed by using BIO-RAD clarity western ECL Substrate.

### 2.5. Yeast mating-based split ubiquitin assay

Split ubiquitin assays were done as described earlier (Obrdlik et al., 2004). The appropriate yeast host strains THY.AP4 and THY.AP5 and gateway cloning-based vectors such as pMetYC gate, pXNgate21-3HA and pXNuB WT gate were used. The TaFLA7 and TaVIH2 templates were PCR amplified using primers containing attB1 and attB2 sites in the forward and reverse primers, respectively. The attB flanking inserts were co-transformed with linearized vectors to the yeast strains using the lithium acetate method for recombination-based invivo cloning. The FLA7 and VIH2 genes were cloned to pMetYC gate and pXNgate 21-3HA vectors, forming C terminal fusions with Cub and NubG, respectively. The Cub vectors and inserts are transformed to haploid THY.AP4 yeast strain and was selected on a synthetic complete medium lacking leucine (SD/-L). Likewise, the Nub vector and inserts are transformed to haploid THY. AP5 yeast strain was selected on a synthetic complete medium lacking tryptophan (SD/-T). The transformed colonies were grown overnight at 28°C in SD media lacking the corresponding dropouts. Subsequently, the cultures were centrifuged and resuspended on a YPDA medium. For interaction studies, mating was performed by mixing an equal amount of the culture, and diploid cells were selected on SD medium lacking the quadruple drop-out of adenine (A), histidine (H), leucine (L), and tryptophan (T). These experiments were repeated three times, and plates were incubated at 28°C for 3-4 days before they were photographed.

### 2.6. Localization studies

TaFLA7 was cloned in pGADT7 vector at EcoR1 and BamH1 sites for the localization experiments. TaVIH2 cDNA was cloned in the pGBKT7 vector using Sma1 and Not1 restriction sites. The constructs were transformed in the Y2H Gold yeast strain and selected on SD-Leu or SD-Trp plates. Yeast spheroplasts were prepared for localization as described earlier (Severance et al., 2004). Mouse monoclonal anti-HA (H.A.:FLA7-pGADT7) : or rabbit Anti-c-Myc (cMYC: VIH2-pGBKT7) primary antibody (Invitrogen, USA) was used for the respective preparations at a ratio of 1:200 followed by 5 washing with blocking buffer. Yeast cells were incubated with Goat Anti-Mouse IgG (H+L) Alexa Flour Plus 488 or Goat Anti-Rabbit IgG (H+L) Alexa Flour Plus 647 (Invitrogen, USA) at a ratio of 1:500 for 4hr at room temperature. Cells were washed with blocking buffer and mounted with Fluor mount (Sigma, USA). Representative fluorescent images were taken using LEICA DM600CS using a 64X oil objective.

### 2.7. Luciferase Assay

Coding sequences (CDS) of TaVIH2 and TaFLA7 were amplified using forward primers containing BamHI and reverse primers containing EcoRI, ensuring that the open reading frame (ORF) remained intact. The amplified products were then cloned into pCAMBIA1300-NLuc and pCAMBIA1300-CLuc vectors. The recombinant plasmids were initially transformed into *E. coli* DH5α cells for propagation and were confirmed by sequencing. Subsequently, the verified plasmids were transformed into the *Agrobacterium tumefaciens* GV3101 strain. Positive clones of GV3101 containing the recombinant plasmids were selected and cultured overnight with appropriate antibiotics. The cells were pelleted and resuspended in freshly prepared induction media containing 10 mM MgCl[, 10 mM MES (pH 5.7), and 150 µM acetosyringone. The optical density (OD) of each culture was adjusted to 0.5, and 1 mL of the Nluc-expressing cells was mixed with 1 mL of the Cluc-expressing cells. The mixture was incubated at 28 °C without shaking for 5–6 hours before agroinfiltration into tobacco leaves (10 µL per spot, with three spots per leaf). The infiltrated plants were maintained in a humidity chamber, kept in the dark for 12 hours, and then exposed to light for 24 hours. Finally, the leaves were harvested, and chemiluminescence was detected, with an exposure time of 20 seconds.

### 2.8. Extraction of Inositol Pyrophosphates

S. cerevisiae cultures were grown at 30 °C in Synthetic complete Dextrose (SD) media until they reached exponential growth, with an A_600_ between 0.6 and 0.8. Cells equivalent to 10 A_600_ units were collected by centrifugation. After washing with ice-cold water, the cells were resuspended in 500 µL of 1 M perchloric acid. The samples were quickly frozen using liquid nitrogen and stored at −80 °C. The samples were briefly vortexed upon thawing, and cell debris was removed by centrifugation at 16,200 g for 5 minutes at 4 °C. The supernatants containing acid-extracted inositol polyphosphates were then purified using titanium dioxide beads (GL Sciences 5020–75000), following the method described by (Wilson and Saiardi 2017). Specifically, the supernatants were combined with 5 mg of TiO_2_ beads per sample, pre-washed with water and 1 M perchloric acid, followed by incubation on a nutator at 4 °C for 20 minutes. The beads with bound inositol polyphosphates were collected by centrifugation at 5,000 g for 1 minute at 4°C, followed by two washes with 1 M perchloric acid. To elute the inositol polyphosphates, the beads were resuspended twice in 250 µL of 3% ammonium hydroxide, rotated for 5 minutes at 4°C, and then centrifuged at 5,000 g for 1 minute. The combined eluates (totalling 500 µL per sample) were dried using a vacuum centrifuge at room temperature for 5 hours. The dried samples were reconstituted in 20 µL of water for capillary electrophoresis electrospray ionization mass spectrometry (CE-MS) analysis.

### 2.9. CE-MS analysis

The analyses were conducted as described earlier (Qiu et al. 2020, 2023; Saiardi et al. 2021). An Agilent CE-QQQ system was used for the analysis, which featured an Agilent 7100 capillary electrophoresis unit, an Agilent 6495C Triple Quadrupole mass spectrometer, and an Agilent Jet Stream electrospray ionization source, all connected via an Agilent CE-ESI-MS interface. The sheath liquid, a 50:50 combination of isopropanol and water, was given at a steady flow rate of 10 µL/min using an isocratic Agilent 1200 LC pump and a splitter to ensure exact flow. A 100 cm long fused silica capillary with an internal diameter of 50 µm and an outside diameter of 365 µm was employed to separate the samples. The background electrolyte (BGE) contained 40 mM ammonium acetate that had been adjusted to pH 9.08 with ammonium hydroxide. Before each sample analysis, the capillary was flushed with BGE for 400 seconds. Samples were delivered into the system by applying a pressure of 100 mbar for 15 seconds, resulting in an injection volume of around 30 nL. The MS source settings included a gas temperature of 150 °C, flow rate of 11 L/min, nebulizer pressure of 8 psi, and sheath gas temperature of 175°C. The capillary voltage was set to -2000V, with a nozzle voltage of 2000 V. Furthermore, the negative high-pressure radio frequency (RF) and negative low-pressure RF were kept at 70 and 40 volts, respectively. The multiple reaction monitoring (MRM) parameters were set up as shown in Table 3. The internal standard (IS) stock solution was created using the described method (Qiu et al. 2020; Saiardi et al. 2021). Which was then added to the samples to help with isomer assignment and quantification of InsPs and PP-InsPs. 5 µL of IS was added to 5 µL of the sample and carefully mixed. IPs were quantified by adding known amounts of the matching heavy isotopic references to the samples. After spiking, the final concentrations in the samples were as follows: 4 μM [^13^C_6_] 2-OH InsP_5_, 20 μM [^13^C_6_] InsP_6_, 1 μM [^13^C_6_] 5-InsP_7_, 1 μM [^13^C_6_] 1-InsP_7_, and 1 µM [^13^C_6_] 1,5-InsP_8_. Dorothea Fiedler provided the [13C6] inositol STD (Harmel et al. 2019).

## 3. Results

### 3.1 VIH proteins contain variable IDR regions

Studies in the yeast and mammalian systems have shown that different inositol pyrophosphate kinase function is based on their protein interaction networks (Gokhale 2013; Rao et al. 2014; Wright and Dyson 2015; Rogers et al. 2021). The human PPIP5K (HsPPIP5K) interaction studies show the typical dual-domains involvement and requirement of the C-terminal intrinsically <u>d</u>isordered region (IDR) for interaction (Machkalyan et al. 2016). This HsPPIP5K contains a large C-terminal IDR region (∼250-550 aa), a missing characteristic for most plant VIHs (Figure 1& Figure S1). The IDR region was checked in the plant VIH proteins. Our analysis indicated that plant VIH proteins have a variable and lower percentage of IDR representation. The plants VIH show 12-22 % of IDR region (Table 1). Interestingly, wheat TaVIH possesses a higher IDR % when compared to the *Arabidopsis* (Figure 1).

**Figure 1:**
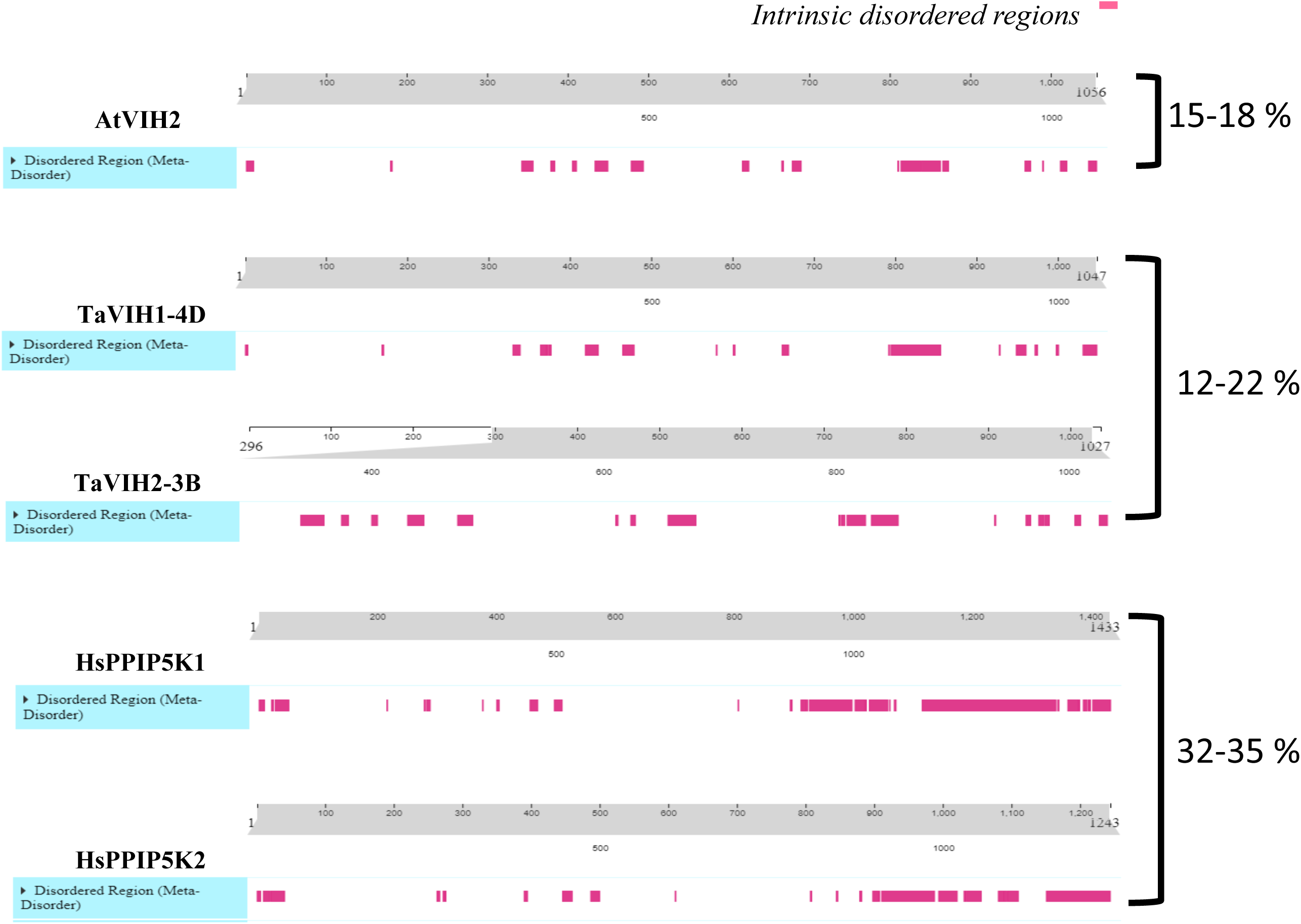
In-silico calculation of the Intrinsic Disorder Regions (IDR) of VIH and PPIP5K proteins. IDR was calculated based on the (Mizianty et al. 2010; Walsh et al. 2012; Han et al. 2023) along with the percentage region score of the mentioned proteins.

**Table 1:**
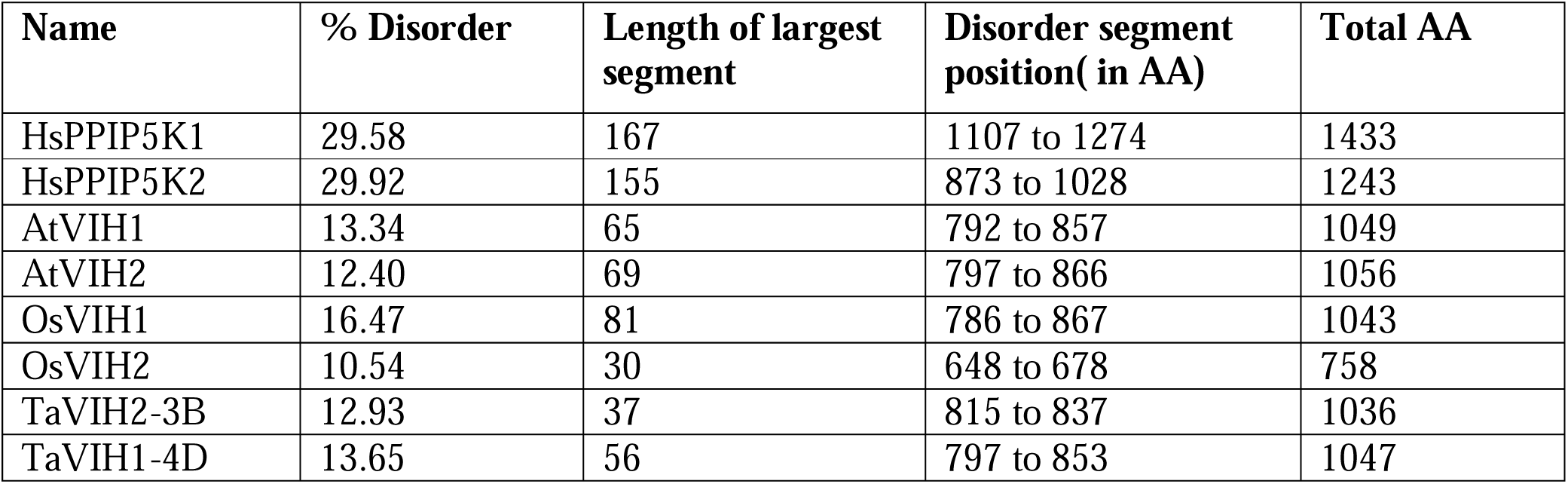
Intrinsic disorder region (IDR) percentage, length and disorder segment position in the human (*Homo sapiens*) and plant (*Arabidospsi thaliana*, At; *Oryza sativa*, Os; *Triticum aestivum*, wheat) encoded PPIP5K-like proteins. Efficient disorder prediction was made using ESpritz (version 1.3).

To get insights into the direct and indirect interactions, we performed a String analysis of VIH proteins. Human PPIP5K and yeast (VIP1) analysis yielded a protein-protein network analysis, with nodes indicating the PHO-mediated interactors for VIP and the involvement of HsPPIP5K in other biochemical activities (Figure S2). A similar analysis of wheat VIH2-3B protein shows multifunctional globular proteins, such as actin, as probable interactors. The presence of IDR in the wheat VIH2 and the preliminary String analysis led us to explore its interacting partners.

### 3.2 Screening of TaVIH2 interacting partners

To get molecular insights into the roleof TaVIH2 proteins in stress tolerance, we used yeast two-hybrid (Y2H) cDNA library screen to identify its possible interacting partners. Protein expression of bait (VIH2) in the yeast cells was confirmed by immunoblot analysis (Supplementary Figure S3A). Two pooled wheat cDNA libraries (∼4 kb to 300 bp) were prepared for Y2H studies. The smear pattern confirmed the good quality of ds-cDNA (Supplementary Figure S3B). Y2H was performed using TaVIH2-3B as a bait, and this resulted in the identification of 89 putative yeast colonies with a mating efficiency of 3.8 %. Subsequent screening of the colonies led to the identification of ∼52 putative interactors (Supplementary Figure S3C&D). Amplicons were generated (Supplementary Figure S4) sequenced, and clones that appeared more than twice were considered for further studies. The analysis led to the shortlisting eleven strong potential interactors with high occurrence in the screening (Table 2). Amongst these interactors, most of the genes encode for cell-wall-related function, including fasciclin-like arabinogalactan protein (FLA), glycosyl-transferases, and glycine-rich structural proteins. The most frequently (nine times) interacting clone was identified as FLA (TraesCS2A02G165600) protein, and further detailed characterization was performed with this protein.

**Table 2:** Interacting partners of wheat VIH2 proteins. The tables indicate the gene IDs with their annotated function based on homology. The numbers indicate the frequency of the occurrence of the clone as an interacting partner from the library.

| S.No | Gene Function | Known functional in Planta | New TRIAE_ ID | Number of Occurrences |
| --- | --- | --- | --- | --- |
| 1. | **Fasciclin-like protein FLA7-2D** | Cell wall biosynthesis, stress response, | TraesCS2D02G172900.1 | 9 |
| 2. | **Glycosyl-transferases (GT1)** | Cell wall biosynthesis, stress response, cell wall cementing, transfer sugar moieties | TraesCS6B02G339100.1 | 6 |
| 3. | **Xylan arabinosyl transferase (Xat1)** | Cell wall biosynthesis, stress response, cell wall cementing | TraesCS6A02G309400.1 | 5 |
| 4. | **Glycine-rich cell wall structural protein-like** | Cell wall biosynthesis, stress response | TraesCS2B02G541900 | 5 |
| 5. | **NADH-ubiquinone reductase complex 1 MLRQ subunit;** | Uncharacterized protein | TraesCS7A02G238600 | 3 |
| 6. | **Hypothetical protein TRIUR3_12806** | Auxin efflux transporter that acts as a negative regulator of light signalling to promote hypocotyl elongation | TraesCS4A02G016700 | 3 |
| 7. | **ABC transporter B family member** | Metal transporter involved in the sequestration of aluminium into vacuoles | TraesCS5A02G392600 | 4 |
| 8. | **Short-chain dehydrogenase reductase** | Glucose signalling and abscisic acid biosynthesis and functions | TraesCS2B02G116700 | 3 |
| 9. | **Ethylene-responsive element binding protein 1 (EREB1)** | Increases tolerance to abiotic stress | TraesCS7B02G062200 | 4 |
| 10 | **W.W. domain-binding protein (containing PPG PPP motif)** | Modules mediate protein-protein interactions by recognizing proline-rich peptide motifs and phosphorylated serine/threonine-proline sites. | TraesCS7B02G062200 | 3 |

**Table 3:** MS parameters for MRM transitions.

| Molecular name | Precursor Ion | Product Ion | Collision Energy (V) | Cell Accelerator Voltage |
| --- | --- | --- | --- | --- |
| [<sup>13</sup>C<sub>6</sub>] InsP<sub>5</sub> | 292 | 504.9 | 9 | 3 |
| InsP<sub>5</sub> | 289 | 498.9 | 9 | 3 |
| [<sup>13</sup>C<sub>6</sub>] InsP<sub>6</sub> | 331.9 | 79.1 | 53 | 4 |
| InsP<sub>6</sub> | 328.9 | 79.1 | 53 | 4 |
| [<sup>13</sup>C<sub>6</sub>] InsP<sub>7</sub> | 371.9 | 322.9 | 9 | 3 |
| [<sup>18</sup>O<sub>2</sub>] 4-InsP<sub>7</sub> | 370.9 | 319.9 | 9 | 3 |
| InsP<sub>7</sub> | 368.9 | 319.9 | 9 | 3 |
| 1,5[<sup>13</sup>C<sub>6</sub>] InsP<sub>8</sub> | 411.9 | 362.8 | 9 | 1 |
| 1,5InsP<sub>8</sub> | 408.9 | 359.8 | 9 | 1 |

Polarity: Negative, Dwell: 80, Fragmentor (V): 166

### 3.3 Wheat VIH2 interacts with fasciclin-like <u>a</u>rabinogalactan protein (FLA)

To further confirm this interaction, full-length coding sequence of TaFLA7 (1.4 kb) was cloned (TaFLA7:AD+TaVIH2:BD) and co-transformed along with TaVIH2 in the yeast strain. No autoactivation of the TaFLA7 and TaVIH2 was observed in the respective AD and BD vectors (Figure S5A). The yeast co-transformed with TaFLA7 and TaVIH2 showed intense reporter activity (Figure 2A). Next, to further confirm this interaction, yeast colonies were grown on -His/-Leu/-Trp (Auerobasidin) media and the protein extracts were subjected to in vivo pull[down assay. For this, first we checked the expression of HA-tagged FLA7 and c-MYC-tagged TaVIH2 in the Y2H Gold yeast strain. Western analysis for both proteins (fused HA:TaFLA7: ∼42 kDa) indicated optimal cell expression (Figure 2B left panel). Co-immunoprecipitation was done using an anti-c-MYC antibody. Furthermore, only the TaVIH2 (∼110 kDa) was detected in the lysate lacking TaFLA7 (Figure 2B right panel; Lane1), whereas no proteins were detected in wild-type yeast Y187 (Lane3). However, in the pull-down analysis of TaFLA7 and TaVIH2 co-transformed cells, both the TaFLA7 and TaVIH2 proteins were detected by respective tagged antibodies (Figure 2C; Lane 4). These findings suggest that TaVIH2 and TaFLA7 exhibit protein-protein interactions. Next, we also employed spilt ubiquitin-based interaction assays to validate these interactions. The respective plasmids were transformed into the respective haploid yeast strains with appropriate constructs, and mating was performed. The yeast strains were streaked on multiple drop-outs, as indicated in Figure 2C. Our results show that yeast containing only TaFL7 or TaVIH2, and their respective controls (1, 2-haploid yeast; 4 and 5-diploid yeast), exhibit no growth. Further specific interactions were noted in diploid yeast strains harbouring both TaFL7 and TaVIH2-containing constructs (3-diploid yeast), suggesting the protein-protein interactions. To further validate these interactions, we performed a split luciferase complementation assay using tobacco leaf (Nicotiana benthamiana) with the fused protein NLuc-TaFLA7)/CLuc-TaVIH2.

**Figure 2:**
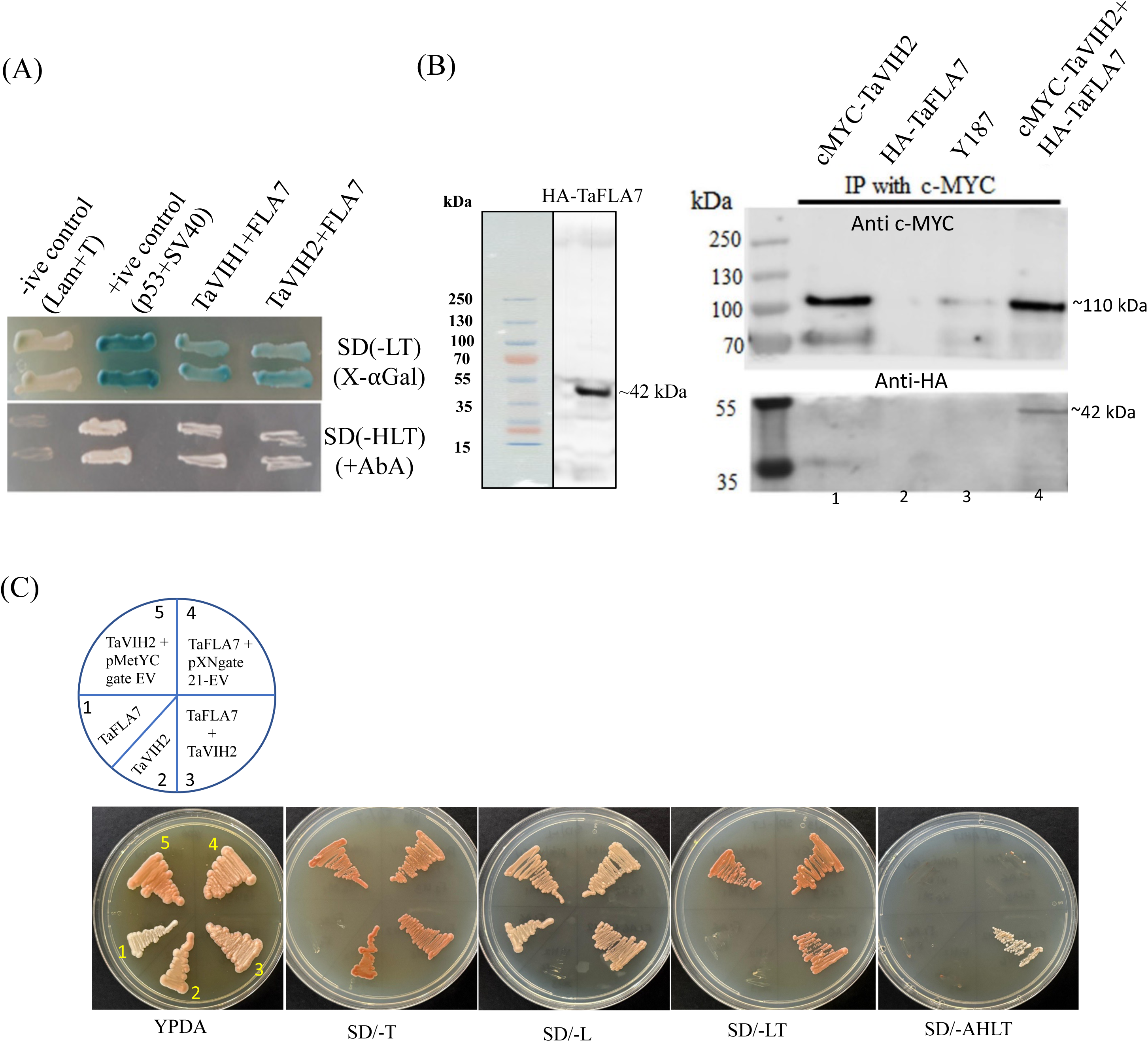
Interaction of TaVIH2-3B with TaFLA7 as identified during Y2H screening. (A) Y2H assay using full-length TaFLA7 and VIH2 as represented by GAL plates containing -L-T and without GAL plates containing -HLT. (C) Western analysis of the wheat FLA7 in the yeast strains harbouring independent clones Lane1: cMYC-TaVIH2; Lane2: HA-TaFLA7; Lane3: Wildtype Y187; Lane4: cotransformed clones of cMYC-TaVIH2+HA-TaFLA7. The numbers indicated the expected molecular weight of the detected proteins. (B) Split-ubiquitin yeast two-hybrid interaction assay of the TaFLA7 and TaVIH2. The appropriate colonies were streaked on single dropout media SD-T (for monokaryon yeast strain with pMETYC:TaVIH2), SD-L for (monokaryon yeast strain pXNgate221:TaFLA7) and double dropout (-LT for dikaryon) and for interactors using quadruple dropout (-AHLT).

Luciferase complementation imaging (LCI) pair reconstituted luciferase activity in regions where both the fused proteins were expressed compared to controls (Figure 3A). RLU measurement suggested an increase of 7-8 folds of the luminescence activity compared to the controls (Figure 3B). These results confirmed that TaVIH2 may interact with proteins involved in cell-wall maintenance.

**Figure 3:**
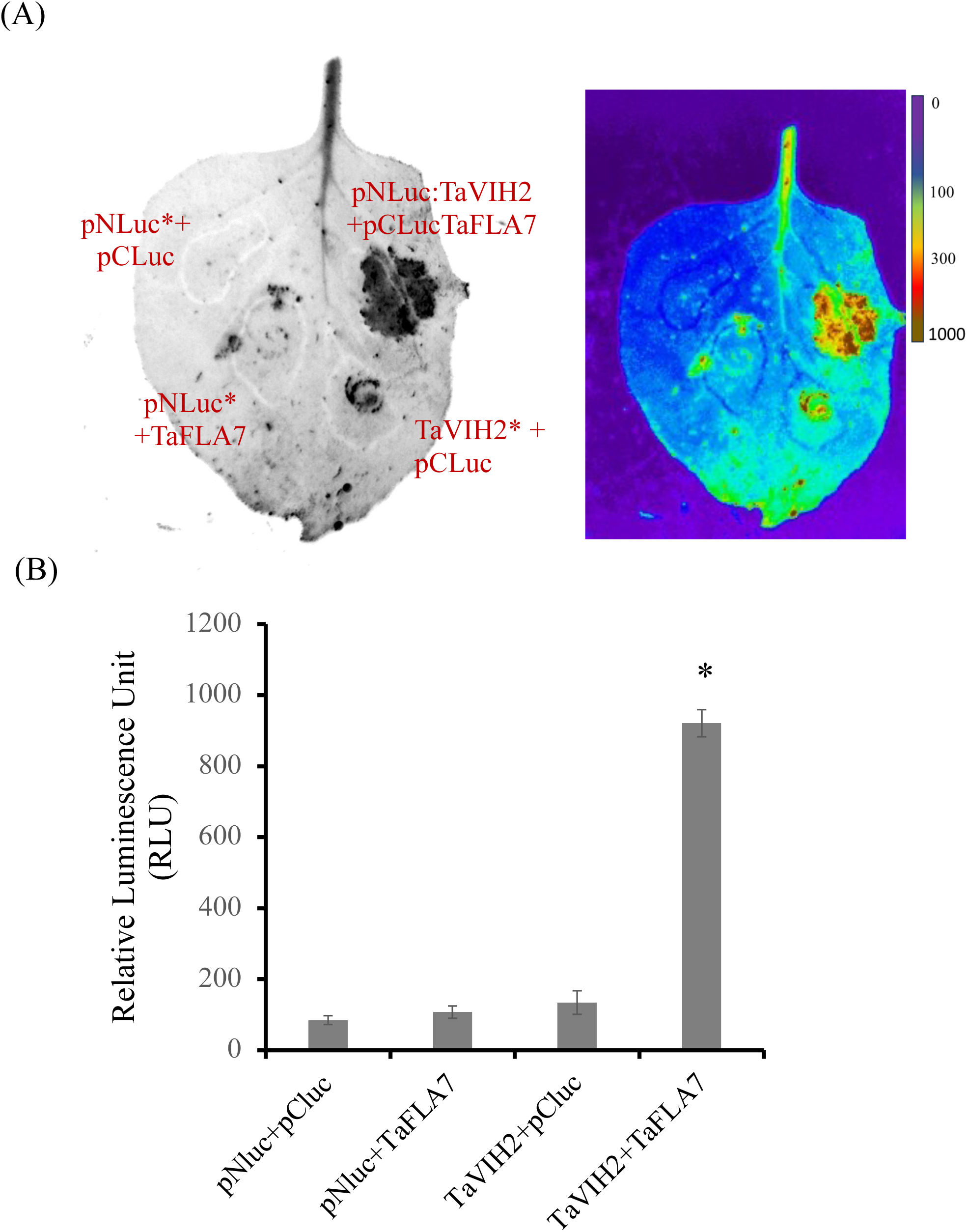
Luciferase reporter assay to characterization of TaFLA7 interaction with VIH2. (A) Luciferase assay of TaFLA7 with VIH2 using tobacco leaves. (B) RLU measurement for the strength of interaction. The values shown are the mean ± SE; n = 4. Student’s t test was performed (*p < 0.001).

### 3.4 VIH2 facilitates the localization of FLA7

Our analysis indicated that TaFLA7 encodes a 367 aa protein (predicted ∼40 kDa) containing FAS-like arabinogalactan protein and the presence of a typical transmembrane domain (TMD) with glycophosphatidyl-inositol (GPI) domain (Figure 4A). The protein hydropathy plot also identified a hydrophobic region near the GPI region at the C-terminal (Figure S5B). Next, we studied the localization of the FLA7 protein in yeast cells. Confocal imaging of yeast cells expressing TaFLA7:GFP suggests its presence on the plasma membrane (Figure 4B). Since the wheat FLA7-VIH2 interact and FLA7 is localized to the membrane, we wonder whether VIH2-3B contributes to the localization of FLA7. Previously, we reported that TaVIH2-3B overexpressing lines enhanced the expression of genes involved in cell wall biosynthesis (Shukla et al. 2021). TaVIH2-3B was shown to utilize InsP_7_ as a substrate likewise to yeast VIP1. Therefore, we tested the localization of FLA7 in different yeast backgrounds. As expected, in the Wild type (WT)( yeast, FLA7 was localized on the plasma membrane but this localisation was affected in the VIP1-defective yeast (Figure 4C). This suggested that localization of the FLA7 was dependent on VIP1 activity. Next, we complemented the yeast vip1Δ mutant and checked for the FLA7 localization. The presence of TaVIH2-3B also restored the localisation of TaFL6 to the plasma membrane in the the yeast vip1Δ strain.

**Figure 4:**
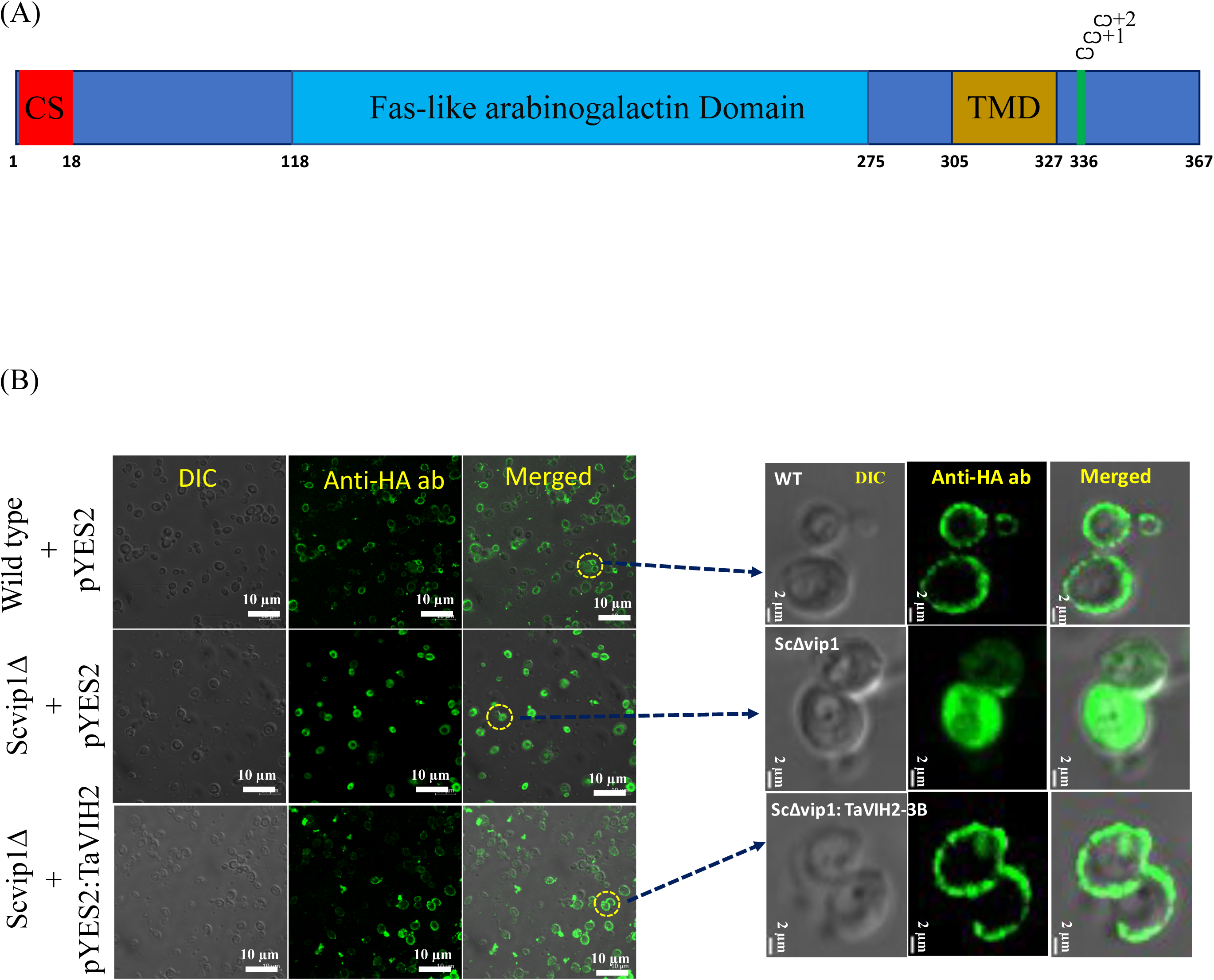
The localization of TaFLA7 in ScVIP1 mutant. (A) Schematic representation of the TaFLA7 protein domain architecture. The N-terminal signal peptide and conserved cysteine-rich (CS) motif are shown in red. The central Fas-like arabinogalactan domain is highlighted in blue, followed by a predicted transmembrane domain (TMD) in yellow. The C-terminal region includes GPI-anchor attachment sites (Ω^1 and Ω^2) predicted by big-PI Plant Predictor. Amino acid positions for each domain are indicated. (B) In situ localization of the TaFLA7 in wild-type and vip1Δ yeast strains. Constructs were introduced under pYES2 vector with or without TaVIH2-3B. HA-tagged TaFLA7 was detected by immunofluorescence. In vip1Δ cells, co-expression with TaVIH2-3B led to increased plasma membrane enrichment of TaFLA7. Right panel: higher magnification images showing plasma membrane enrichment in wild-type and vip1Δ cells. Scale bars are indicated for each panel. Fluorescence was measured using a Zeiss fluorescence microscope (Axio Imager Z2) with an Axiocam MRm camera at 63X. Representative images are shown and similar observations were noted for 3-4 independent transformed colonies of yeast.

Additionally, we have also profiled the cellular pools of InsP_6_, 5-InsP_7_, 1-InsP_7_, and 1,5-InsP_8_ with capillary electrophoresis electrospray ionization mass spectrometry (CE-ESI-MS), as described in (Qiu et al. 2020, 2023; Saiardi et al. 2021; Liu et al. 2023; Eisenbeis et al. 2023). This method allows us to investigate the cellular levels of different InsP and PP-InsP species in the yeast transformants. To account for any differences in sample recovery during extraction and TiO_2_ bead affinity enrichment, the levels of 5-InsP_7_, 1-InsP_7_, and 1,5-InsP_8_ were calculated and plotted in pMol (Figure 5). The observed changes in the 1,5-InsP_8_ levels (Figure 5) between the WT, vip1Δ mutant and vip1Δ::TaVIH2-3B complemented extracts confirmed that TaVIH2 is a functional InsP_8_ synthase in yeast and it is likely that the localization of TaFLA7 is dependent of the TaVIH2-derived InsP_8_. Collectively, our results indicate that TaVIH2-3B interacts with the FLA7 protein and could facilitate its localization to the plasma membrane. Moreover, the differences in 1,5-InsP_8_ levels witnessed among the WT, *vip1*Δ mutant, and *vip1*Δ mutant complemented with TaVIH2-3B suggest that the appropriate positioning of TaFLA7 could be associated with the activity of the PPIP5K enzyme.

**Figure 5:**
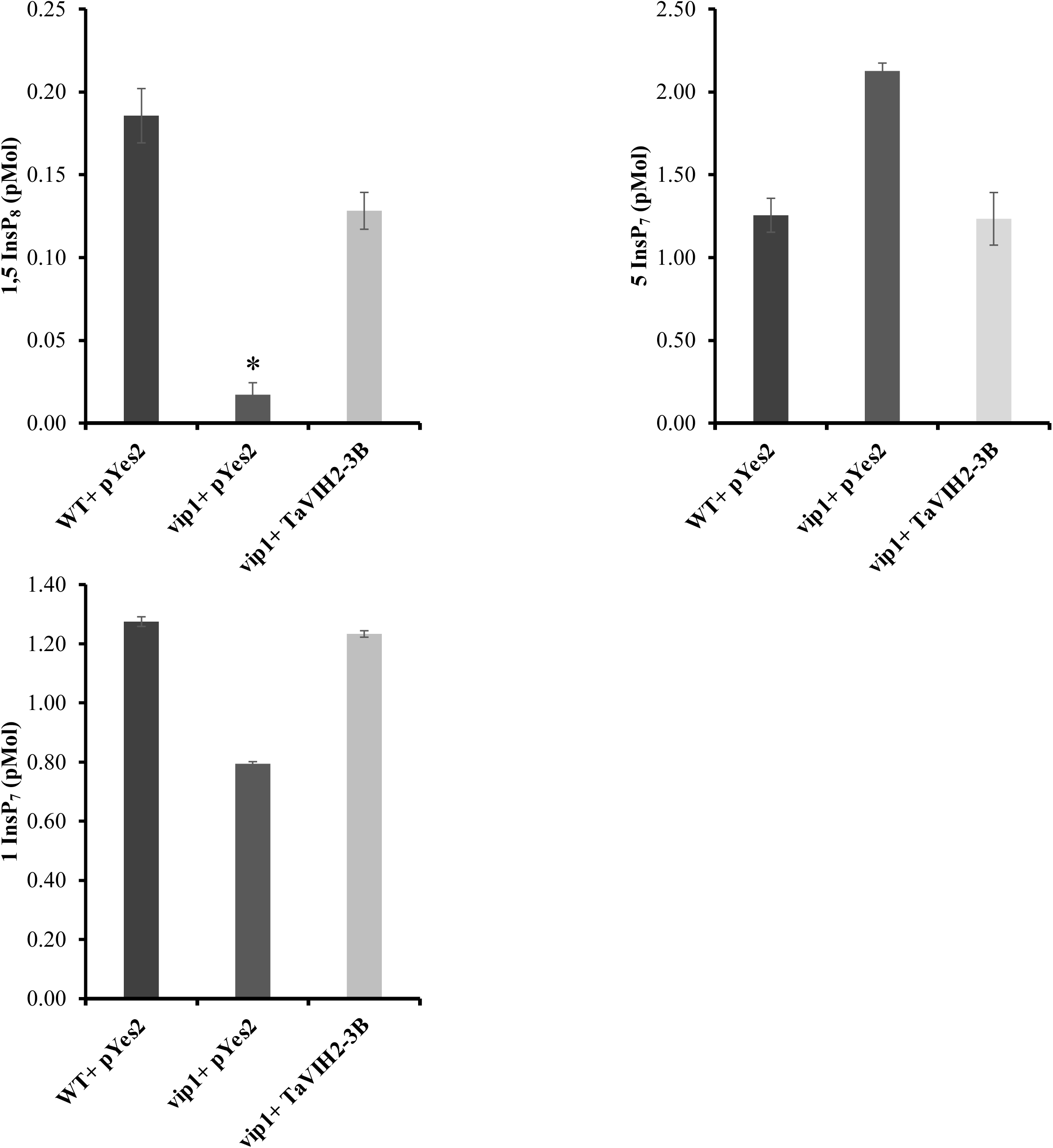
Quantification of inositol pyrophosphate levels using CE-ESI-MS. The inositol pyrophosphate levels in extracts of the WT + pYES2, vip1Δ + pYES2, and vip1Δ + TaVIH2-3B strains that were transformed with TaFLA7 are displayed for 1,5-InsP_8_, 5-InsP_7_, and 1-InsP_7_. The values shown are the mean ± SE; n = 5. Student’s t test was performed (*p < 0.001).

To investigate whether FLA proteins are the possible cellular targets of TaVIH-derived InsP species, we performed in silico docking experiment using the structural model of wheat FLA7 closest homoeolog viz. TaFLA7-2D and TaFLA7-2A (TraesCS2A02G165600) with various InsP isomers. Notably, the InsP_6_ was docked robustly at the N-terminal domain of TaFLA7-2A and TaFLA7-2D proteins (Figure 6). Furthermore, docking with 5-InsP_7_ and 1,5-InsP_8_, the putative products of TaVIH, revealed that the binding pocket remained unchanged, with key conserved residues contributing to ligand interactions. In TaFLA7-2A, Ser239, Thr256, and His253 were consistently involved in binding all three ligands, whereas in TaFLA7, His228, Ser245, and Asn275 were coordinating with the InsP isomers. This highlights the potential of these FLA proteins to accommodate diverse inositol phosphate ligands. Additionally, truncated versions of the proteins, lacking the transmembrane (TMD) domain, retained identical binding pockets and conserved residues, indicating that the TMD does not influence ligand binding (Figure S6). These findings support the structural and functional integrity of the ligand-binding domain in TaFLA7, independent of the TMD.

**Figure 6:**
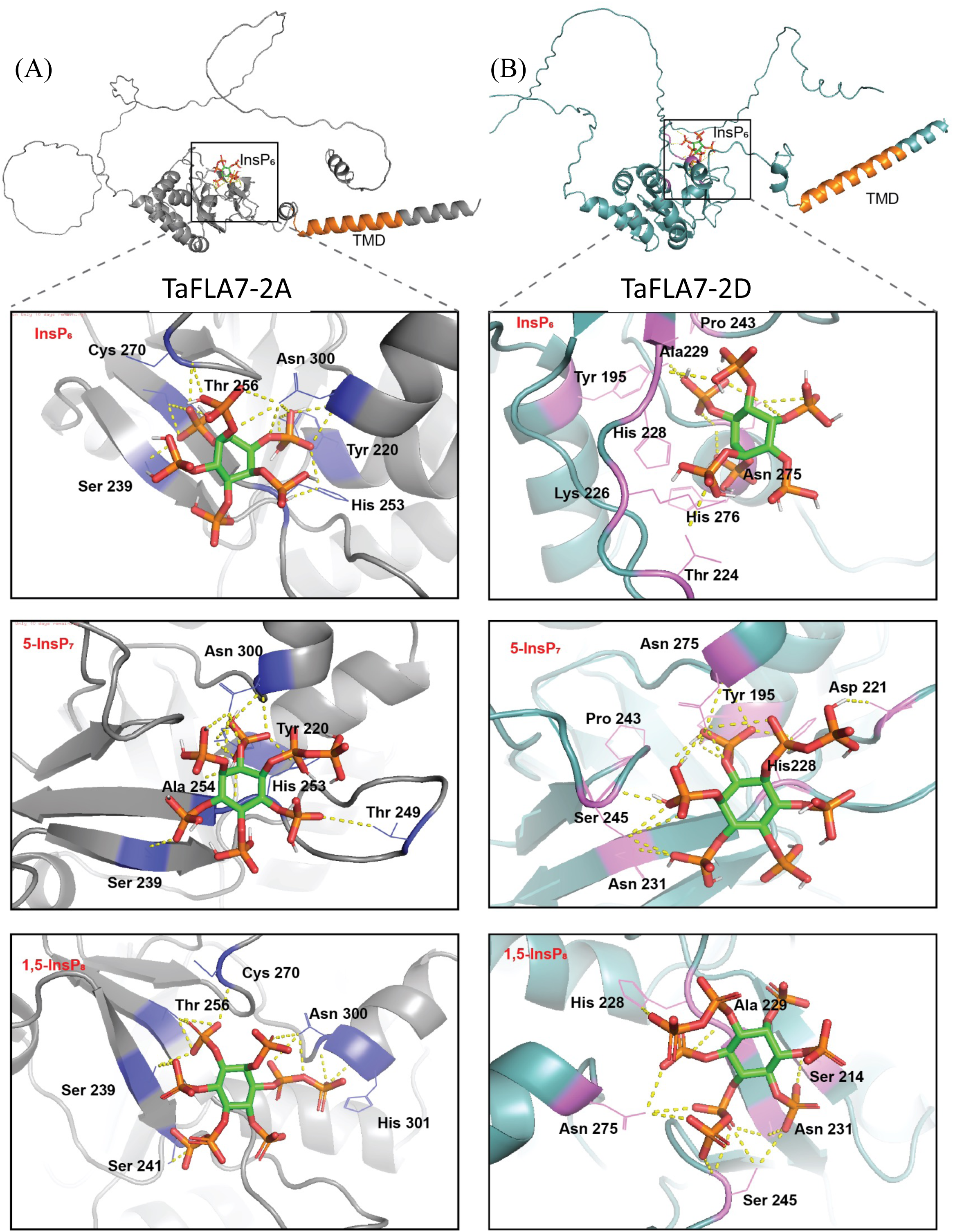
Molecular docking of TaFLA7 homoeologs (TaFLA7-2A & 2D) with InsP_6_, 5-InsP_7_, and 1,5-InsP_8_. Schematic representation of (A) TaFLA7-2A and (B) TaFLA7-2D structures, highlighting the transmembrane domain (TMD) and ligand-binding site. Docked conformations of InsP_6_, 5-InsP_7_, and 1,5-InsP_8_ are shown with key interacting residues labelled. Ligands (green sticks) form polar bonds (yellow dashed lines), revealing distinct binding orientations and interaction patterns.

### 3.5. FLA7 is differentially expressed in wheat tissue

The FLA proteins are known to respond to different stress conditions and are involved in plant growth and development (Zang et al. 2015; Cagnola et al. 2018; Wu et al. 2020; Ma et al. 2022; Yao et al. 2023). These observations suggest that VIH2 and its interacting partners could participate in similar pathways. To further characterize the wheat FLA7, we studied its expression in different tissues and under various stress conditions. The *TaFLA7* was expressed in the grain at the Zadoks scale (Z) 71 stage (watery ripe with the start of grain filling) and in the spike at Z65 (50% of the spikes had yellow anthers) of wheat development (Figure S7A). The transcript expression of *TaFLA7* showed significant upregulation in the shoot tissue when subjected to drought stress after 1 hr and 6 hrs (Figure S7B). Our quantitative RT-PCR results suggested a significant upregulation in the shoot or root tissue subjected to drought stress (Figure S7C). Quantitative RT-PCR showed a high expression of *TaFLA7* towards 28 days after the anthesis (DAA) stages of grain development (Figure S7D). Expression analysis of *TaFLA7* in different wheat grain tissue using the eFP wheat browser revealed its high expression in embryo and aleurone layers of the grain (Figure S8). The tissue-specific eFP wheat browser analysis indicates the high expression of *TaFLA7* in grains, shoot axis, and spikelets (Figure S9). Altogether, the expression data of the *TaFLA7* show transcriptional changes during desiccation, a prerequisite step in grain maturation.

## 4. Discussion

Protein-protein interaction networks provide valuable insights into biological processes and complex metabolic functions within living cells. The Y2H system leverages eukaryotic transcriptional activators with distinct functional domains for DNA-binding and transactivation, making it a powerful and rapid approach to discovering new protein interactions (He et al. 2019). In this study, a high-quality cDNA library using RNA extracted from wheat seedlings was constructed. Our work identified several interactors of TaVIH2 proteins, which serve as promising entry points for further exploration of their regulatory network in the developmental process. Recently, it was demonstrated that the Triticum aestivum (hexaploid wheat) genome encodes functional PPIP5K isoforms, identified as TaVIH2-3B (Shukla et al. 2021). Moreover, the heterologous expression of wheat TaVIH2-3B in Arabidopsis imparts drought tolerance. These Arabidopsis transgenic lines expressing TaVIH2 also exhibit a reprogramming of gene expression involved in cell-wall metabolism and stress response pathways (Shukla et al. 2021). In particular differential expression of genes linked to ABA biosynthesis, cell wall maintenance, and cytochrome P-450 monooxygenases were highly upregulated. Although the level of InsPs seems to be important in imparting such a function, it remains to be studied if plant PP-InsP synthase, such as VIH at the protein level, could be involved directly.

Multi-protein interactions were more accessible by IDRs, which helped them connect with other proteins in the cell. Abelson (Abl), a protein non-receptor tyrosine kinase, has an IDR domain (around 900 amino acids) that has been implicated in a variety of tasks in Drosophila, including the regulation of cell shape and cytoskeleton development (Rogers et al. 2021). Similarly, the human PPIP5K domains’ association with proteins involved in actin cytoskeleton structure and phosphatidylinositol biosynthesis was attributed to the IDR’s existence (Machkalyan et al. 2016). The interaction studies were pursued using human PPIP5K (HsPPIP5K) (Machkalyan et al. 2016). Human PPIP5K, besides containing the typical dual-domain, also possesses a C-terminal IDR (Machkalyan et al. 2016). This IDR in HsPPIP5K spans a large portion (∼250-550 amino acids), a feature that is not very common in most plant VIHs (Figure S1). IDRs assist in their interaction with multiple proteins to facilitate their cellular activities (Ward et al. 2004).

Although VIH proteins have been identified in several plants (Laha et al. 2015; Zhu et al. 2019), including wheat (Shukla et al. 2021), the IDR domains are relatively smaller, ranging from approximately 33-56 amino acids, and their functional roles remain to be elucidated. Our network analysis using the String database can provide clues for the regulatory network of the VIH proteins (Figure S2). Human PPIP5K and yeast (VIP1) analysis resulted in the protein-protein network analysis with nodes pointing towards the PHO-mediated interactors for the VIP and involvement of HsPPIP5K in other biochemical activities (Figure S2). A similar analysis of wheat VIH shows a network with cytoskeletal actin as a probable interactor. Actin is involved in forming microfilaments in the cytoskeleton and can modulate the cellular shape (Pollard and Cooper 2009). This study identified multiple interactors of VIH2, yet most of them are speculated to be involved in cell wall maintenance. The speculative parallels of VIP1 in yeast and VIH in lets us postulate that plant VIH is likely involved in cell wall cementing and homeostasis. The conserved role of VIH was demonstrated in the important cellular function of Pi-homeostasis (Dong et al. 2019; Zhu et al. 2019; Riemer et al. 2021). Under nutrient-limiting conditions, VIP1 and its ortholog in Schizosaccharomyces pombe (Asp1) kinase were shown to control physiological changes such as dimorphic switch, cell-to-cell, cell-substrate adhesion and unipolar budding (Pöhlmann and Fleig 2010). The ability of TaVIH2 to facilitate the localization of TaFLA7 (Figure 4) provides new insight into the functioning of VIH proteins. Based on our observations and previous reports, the involvement of VIH proteins points towards their role in modulating the cellular boundaries and shape (Figure 7). Such phenotypic changes, including the formation of pseudo-hyphal filaments in yeast (Norman et al. 2018), help sustain themselves under stress conditions, yet their similar involvement in plants needs more investigation. Previous biochemical investigations of TaVIH2-3B overexpressing lines in Arabidopsis indicated changes in cell wall cementing metabolites such as cellulose, arabinoxylan, and arabino-galactan (Shukla et al. 2021). Furthermore, RNAseq analysis of the Col-0 and TaVIH2 overexpressing lines suggested high expression of genes involved in cell wall biosynthesis (Shukla et al. 2021). Together with these results, our findings suggest that VIH2 can regulate the composition of cell wall cementing pathways by localizing the involved proteins to their correct positions. The intriguing role of VIH protein overexpression in modulating cell wall composition requires further investigation. Specifically, the in-vivo levels of InsP_7_/InsP_8_ (Figure 5) confirm that the PPIP5K enzymatic activity previously demonstrated with the purified TaVIH2-3B kinase (Shukla et al., 2021) is also functionally active in yeast cells. Therefore, either the synthesis of InsP_8_ or the presence of TaVIH2-3B protein in the yeast cell are directly influencing the correct localization of the TaFLA7 to the plasma membrane. These measurements therefore validate that TaVIH2-3B maintains its PPIP5K activity in vivo, as evidenced by its ability to modulate InsP_7_/InsP_8_ levels in a heterologous yeast system. This, in turn, supports its proposed role in regulating interacting partners such as TaFLA7.

**Figure 7:**
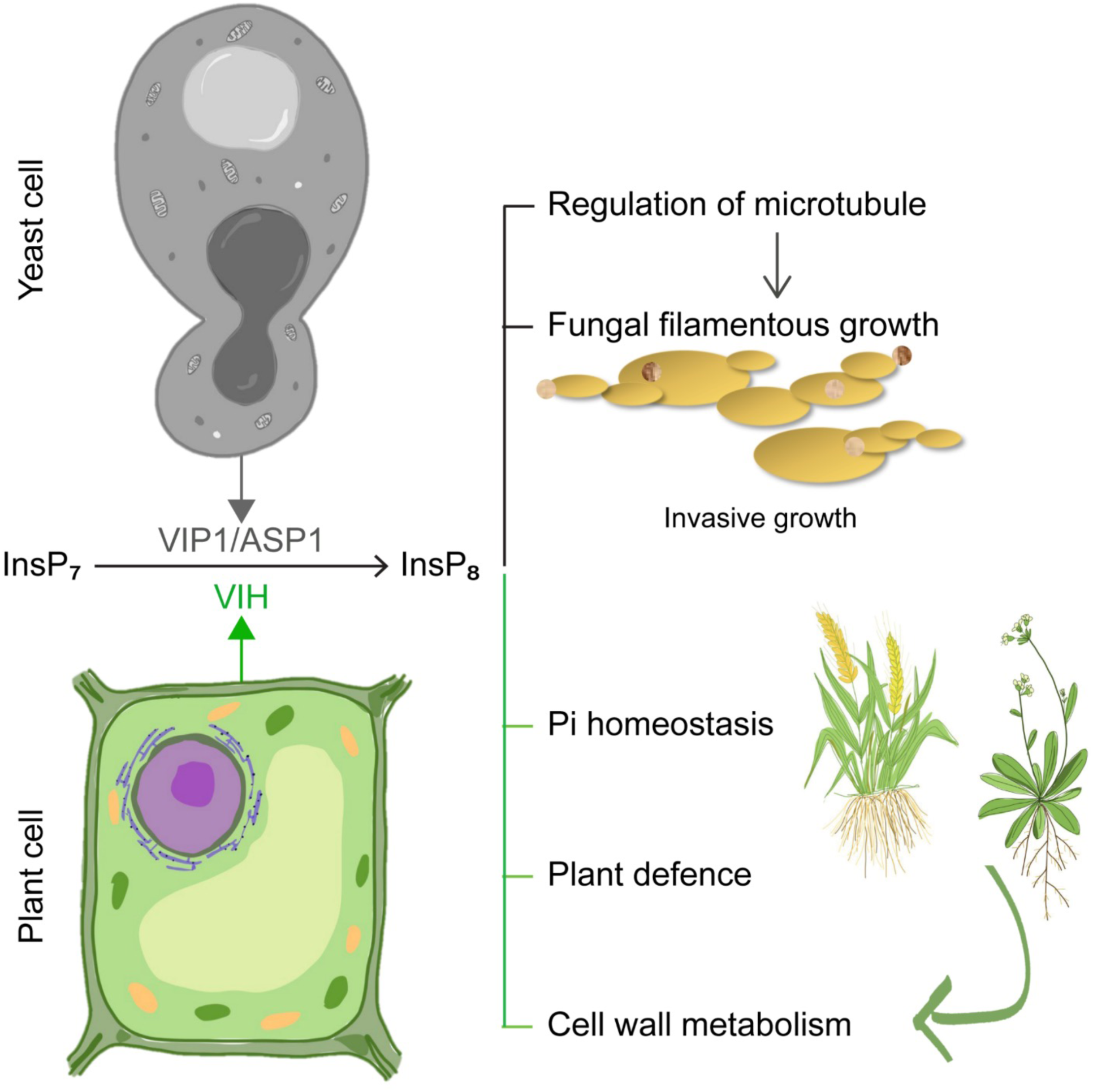
Schematic representation drawing parallels for the possible function of VIH proteins in multiple cellular processes. In yeast, VIP proteins and their products were shown to be involved in the cell boundary rearrangements, including regulation of microtubule rearrangement and filamentous growth of the fungi under nutrient-limiting conditions (Pöhlmann and Fleig 2010; Pöhlmann et al. 2014). Plant VIHs are known to be involved in phosphate (Pi) homeostasis and plant defence (Laha et al. 2015; Dong et al. 2019; Riemer et al. 2021). Recently, the role of plant VIH in cell wall-related rearrangements has been established (Shukla et al. 2021). The participation of genes encoding VIP1/VIH proteins in cell wall rearrangement in both yeast and plants could reflect the conserved function of the proteins across species (Pöhlmann et al. 2014; Shukla et al. 2021).

Furthermore, it is crucial to investigate how this localization occurs and to identify which region of TaFLA7 interacts with TaVIH2-3B and understanding the role in such interaction may also be critical for future study. In the aftermath of the current study, it will be interesting to check if InsP_7_/InsP_8_ could modulate the protein-protein interactions of transcription factors that are primarily involved in water-deficient stress conditions. Although as primary evidence, our docking study provides insight into the ability of InsPx to bind specific regions of FLA7 (Figure S6). This study remains to be further investigated so as to unravel the new mechanistic insights into drought tolerance that could be mediated by cell wall maintenance. Studies using Arabidopsis have deciphered multiple roles of VIHs in different biological processes. However, such efforts should be expanded to crop plants such as wheat for translational research.

## Acknowledgements

The authors thank the Executive Director for the facilities and support. This study was supported by the Department of Biotechnology, Basic Plant Biology Grant to AKP [BT/PR12432/BPA/118/35/2014]. Part of this work was also supported by the NABI-CORE grant to A.K.P. and by the Deutsche Forschungsgemeinschaft (DFG) under Germany’s Excellence Strategy CIBSS, EXC-2189, Project ID 390939984 to H.J. DBT-eLibrary Consortium (DeLCON) is acknowledged for providing timely support and access to e-resources for this work. A.S. acknowledges Centre For Integrative Biological Signalling Studies for PostDoctral Fellowship. D. L. acknowledges the Department of Biotechnology (DBT) for HGK-Innovative Young Biotechnologist Award (BT/13/IYBA/2020/04). We thank Dr. Santosh Satbhai (IISER-M) for sharing reagents for Yeast mating-based Split Ubiquitin Assay.

## Declarations

### Data Availability

No new datasets were generated during the current study.

