## SupplementaryTables for "The wheat VIH2-3B, a functional PPIP5K controls the localization of fasciclin-like arabinogalactan protein"

**Table1: Primers used in the study**

| **Gene** | **Nucleotide Sequence (5'-3')** |
| --- | --- |
| *TaFLA7*  *(full length cloning)* | F: GATGGCCGCACCTTTTCTGCTCCTTCTTCTGCTCGTC |
|  | R: TTAACCAGATAAATTGGCCTTGTTGTTCATCTTGTC |
| *TaFLA7 (subcloning)* | F: GAGAGGGATCCATGGCCGCACCTTTTCTGCTCG |
|  | R: CTCTAGAATTCTTAACCAGATAAATTGGCCTTGTT |
| *BD amplimer primers* | F: GGGAATTCCATATGATGGCCGCACCTTTTCTGCTCGTTCTTCTGCTCGTC |
|  | R: CTCTAGAATTCTTAACCAGATAAATTGGCCTTGTT |
| *TaFLA7 qRT-PCR* | F: ACGACTCAGGCCGTGGGCACGCGT |
|  | R: CGTCCGTGAACGCCCGCTCCTCC |
| *TaARF qRT-PCR* | F: TGATAGGGAACGTGTTGTTGAGGC |
|  | R: AGCCAGTCAAGACCCTCGTACAAC |

**Table 2: Capillary electrophoresis-mass spectrometry (CE-MS) analysis of the different InsPs**. The InsPs were measured in TAFLA7 transformed wild-type yeast (BY4741:pYES2), vip1 mutant (vip1: pYES2) and vip1∆ complemented with TaVIH2-3B (vip1: TaVIH2-3B). The concentrations measured were in picomoles form 3 independent replicates.

|  | **Conc.(pmol)** | | | | |
| --- | --- | --- | --- | --- | --- |
|  | **InsP_5_** | **InsP_6_** | **1 InsP_7_** | **5 InsP_7_** | **1, 5 InsP_8_** |
| **4741 + pYES2 a** | 4.667 | 15.689 | 1.465 | 1.216 | 0.204 |
| **4741 + pYES2 b** | 5.638 | 16.644 | 1.199 | 1.371 | 0.173 |
| **4741 + pYES2c** | 5.222 | 17.192 | 1.161 | 1.178 | 0.180 |
| **vip1 + pYES2a** | 5.857 | 15.734 | 0.632 | 2.139 | 0.015 |
| **vip1 + pYES2b** | 5.205 | 16.516 | 0.846 | 2.166 | 0.025 |
| **vip1 + pYES2c** | 4.473 | 21.342 | 0.905 | 2.074 | 0.011 |
| **vip1 + (TaVIH2-3B) a** | 8.485 | 17.696 | 1.290 | 1.072 | 0.137 |
| **vip1 + (TaVIH2-3B) b** | 7.700 | 25.412 | 1.249 | 1.240 | 0.116 |
| **vip1 + (TaVIH2-3B) c** | 5.194 | 13.923 | 1.160 | 1.389 | 0.131 |
