## SupplementaryFigures for "The wheat VIH2-3B, a functional PPIP5K controls the localization of fasciclin-like arabinogalactan protein"

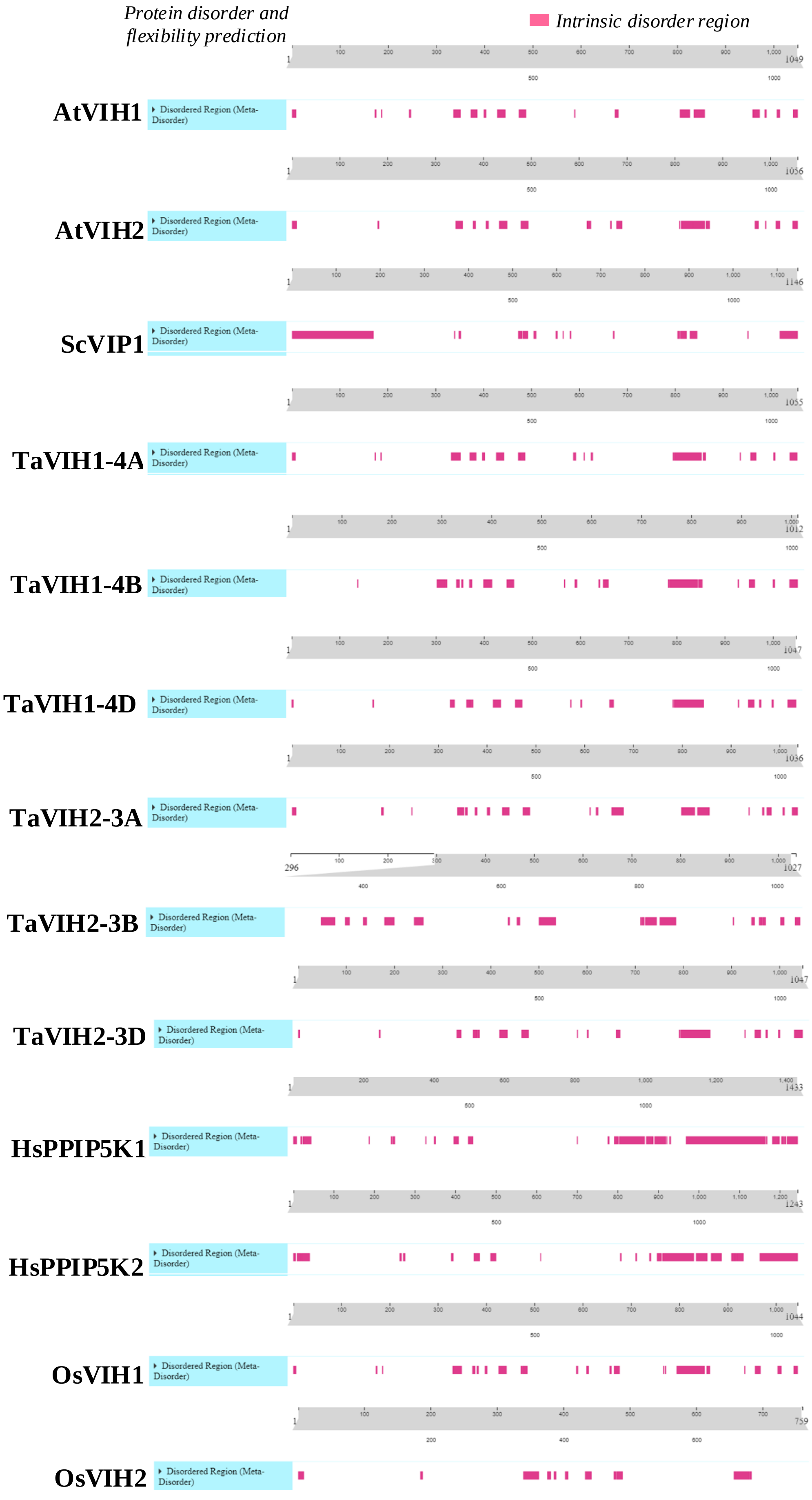


**Supplementary Figure S1:** IDR analysis of VIH/VIP protein sequences. The orange region indicates the predicted IDR regions**.**

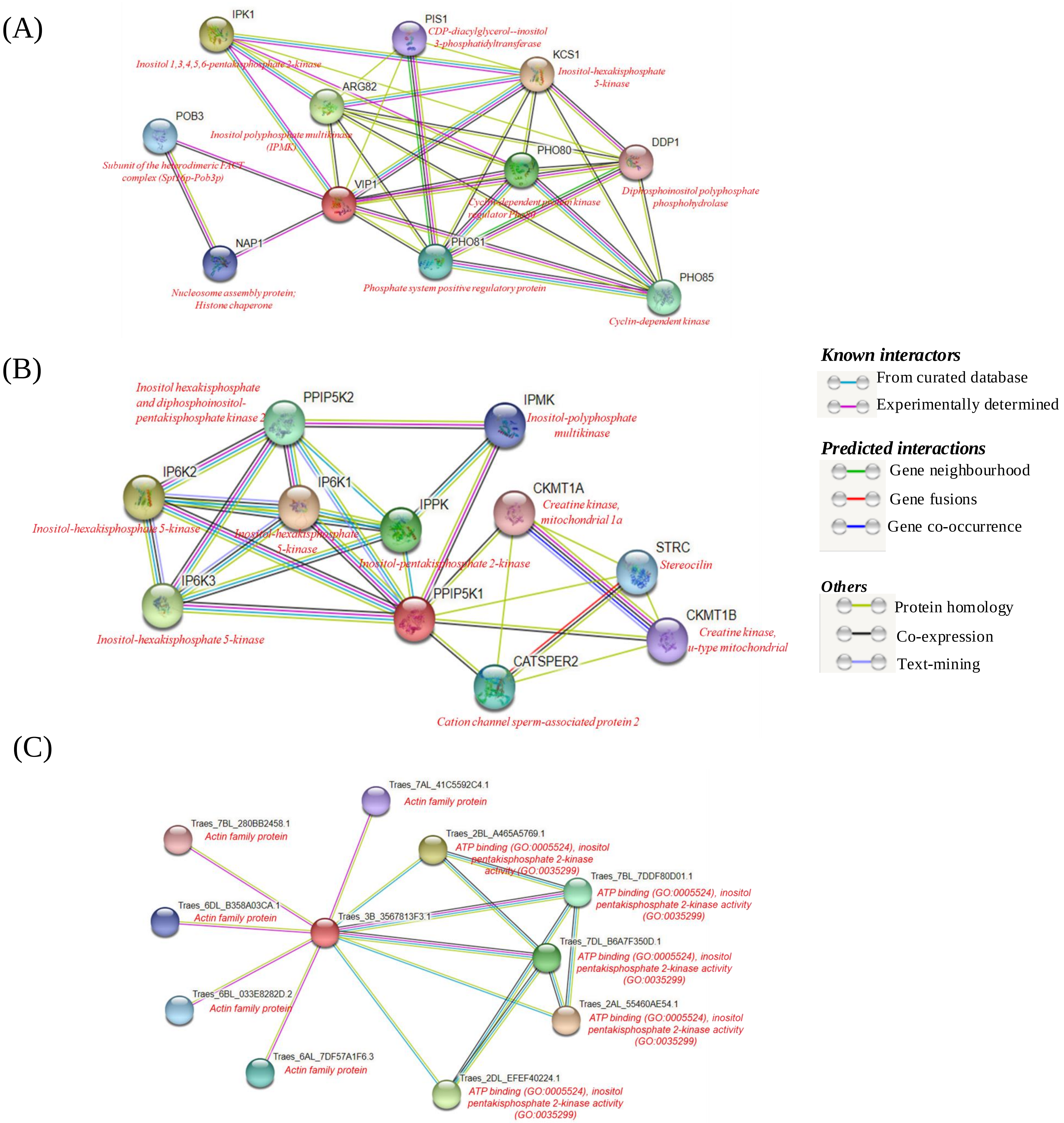


**Supplementary Figure S2:** Protein association networks were identified for each of the human (PPIP5K1), yeast (VIP1) and wheat (VIH; Traes_3B_3567813F3.1). The network analysis for each of the proteins was performed using STRING and was represented as the coloured lines with the description as given in the figure.

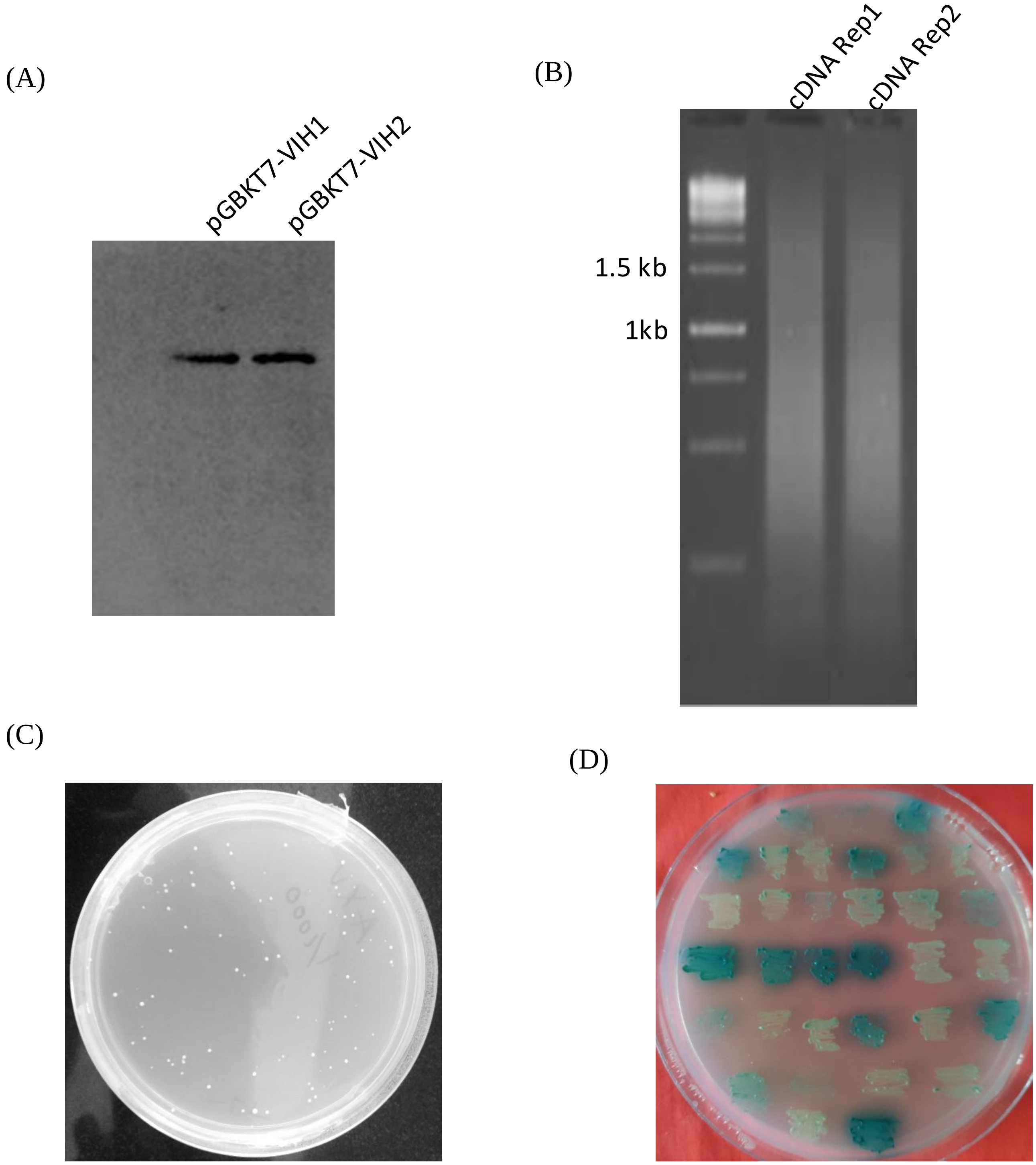


**Supplementary Figure S3:** Western blot analysis, cDNA preparation and screening of the VIH interacting proteins (representative image). (A) Western analysis of c-MYC fused TaVIH1 or TaVIH2 proteins in the yeast strain. (B) Double strand cDNA preparation of wheat seedling library used for yeast two hybrid interaction studies. The cDNA library was resolved on the 1.2 % agarose gel. Two different cDNA preparations (Rep1 and Rep2) was performed and pooled together for the library preparation. (C) Representative yeast colony map for calculating the mating efficiency (-LT$_{1/1000}$). (D) Representative picture of the yeast colonies with putative interacting clones obtained on the selection plates (-AHLT+αGal). Each of the independent streaked colonies represent single putative interacting clone. Only colonies showing strong-blue coloration were used further for further study.

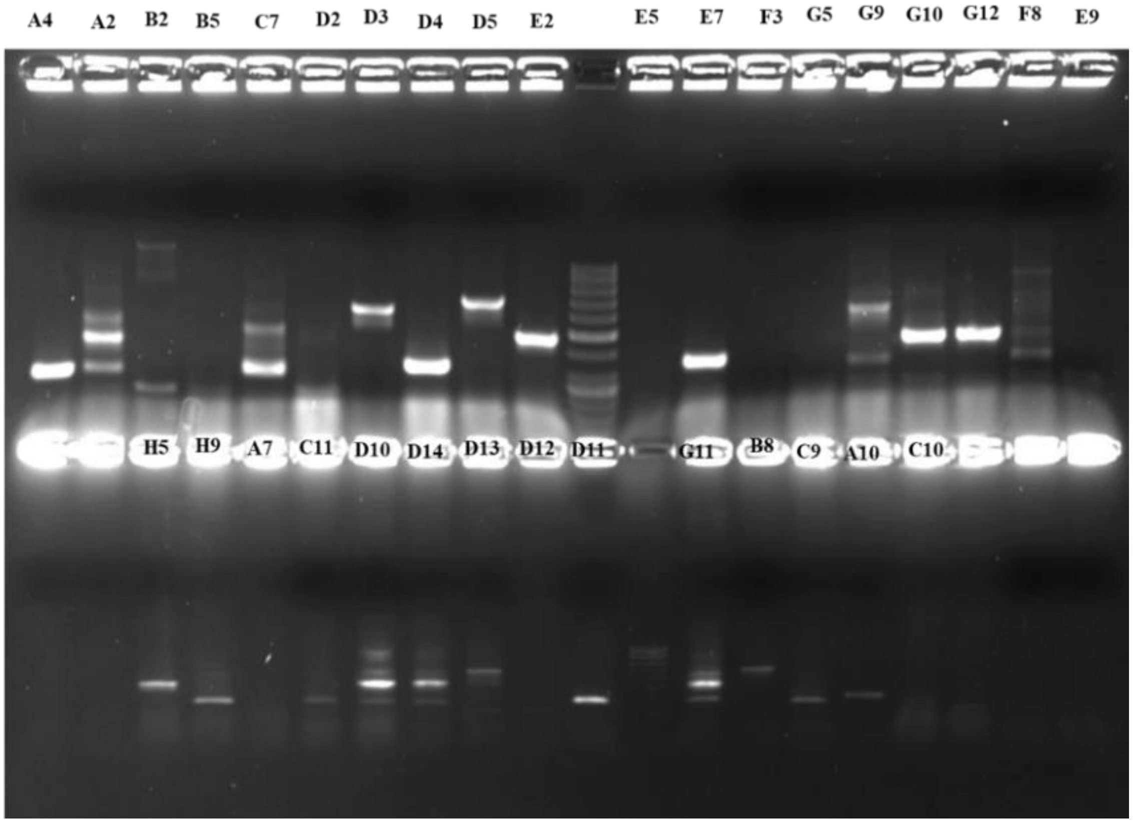


**Supplementary Figure S4:** Representative picture for the colony PCR analysis of yeast blue colonies to identify the interacting partners. The amplicons were cloned in pGEMT-Easy vector and sequencing was performed. The numbers on the lane represent independent colonies of yeast used for amplification of the cDNA clones.

(A)

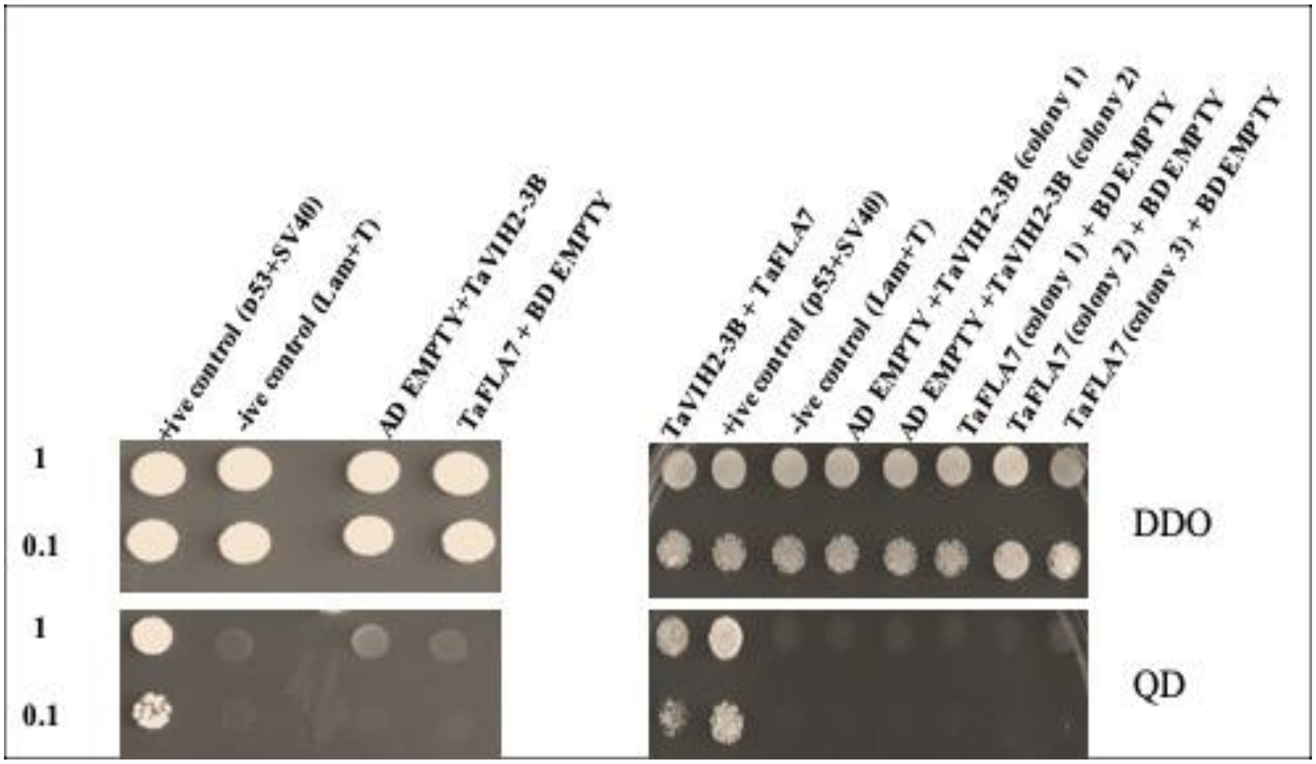


(B)

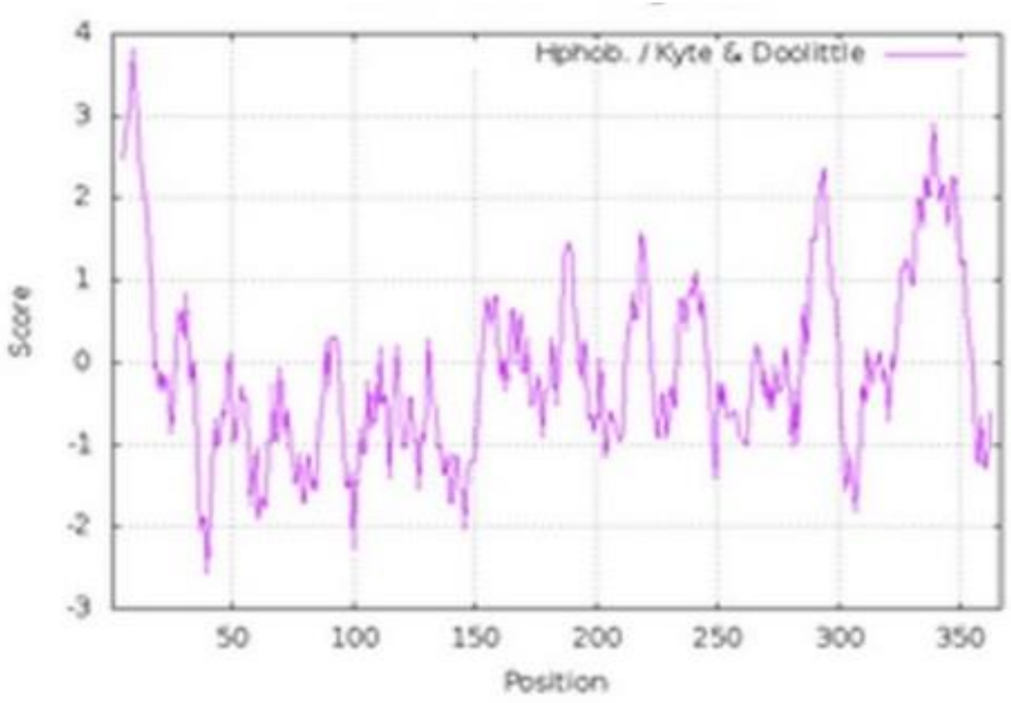


**Supplementary Figure S5: (A)** Autoactivation test for the taVIH2 and FLA7 clones in double dropout (DDO) and quadruple drop out (QD). (B) Hydropathy plot for TaFLA7 domains. The negative values on the Y-axis indicates the degree score of the hydrophilic regions.

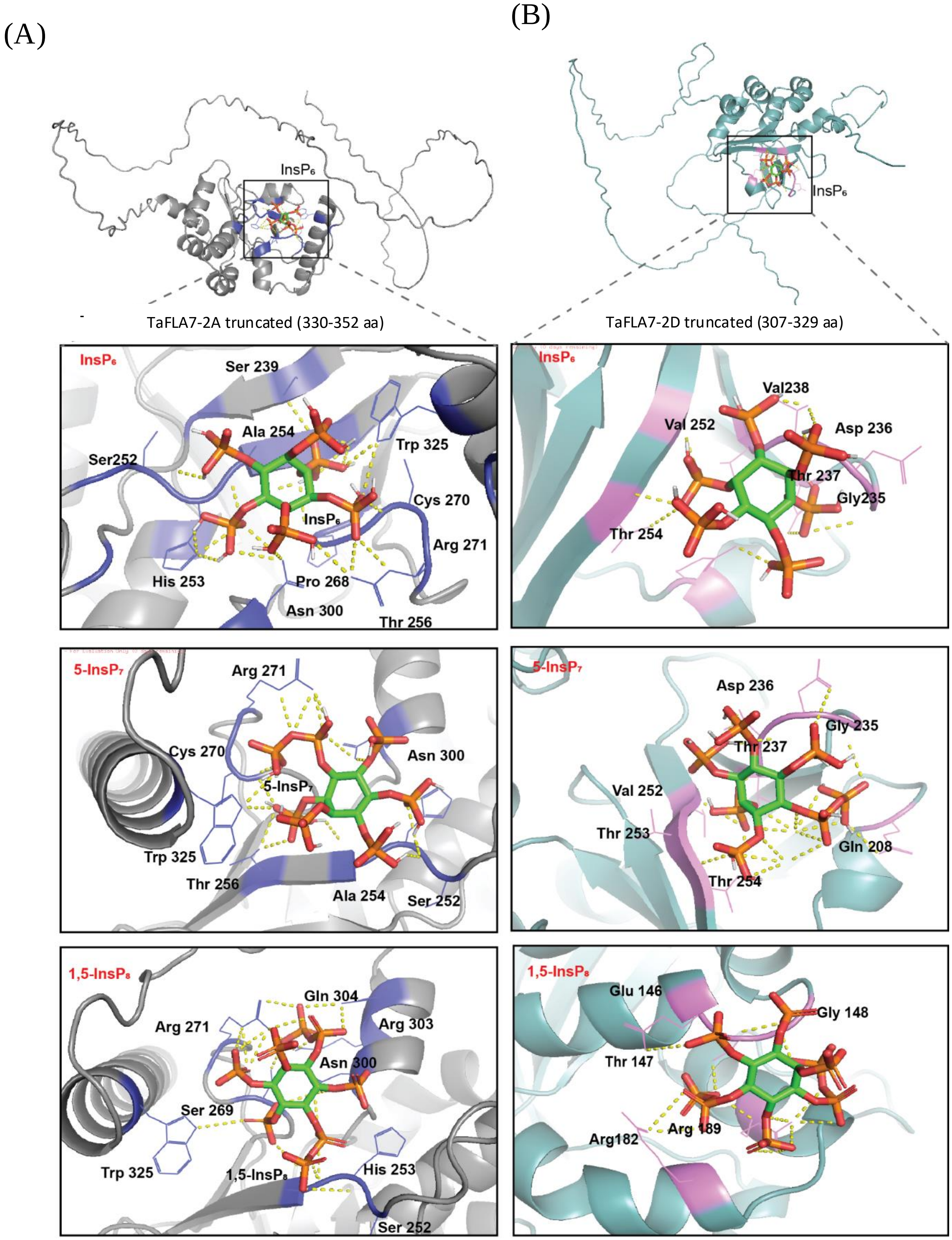


**Supplementary Figure S6:** Molecular docking of truncated (TMD absent) TaFLA7-2A and TaFLA7-2D with $InsP_6$, 5-$InsP_7$, and 1,5-$InsP_8$. Schematic representation of (A) TaFLA7-2A truncated and (B) TaFLA7-2D truncated structures. Docked conformations of $InsP_6$, 5-$InsP_7$, and 1,5-$InsP_8$ are shown with key interacting residues labelled. Ligands (green sticks) form polar bonds (yellow dashed lines), revealing distinct binding orientations and interaction patterns.

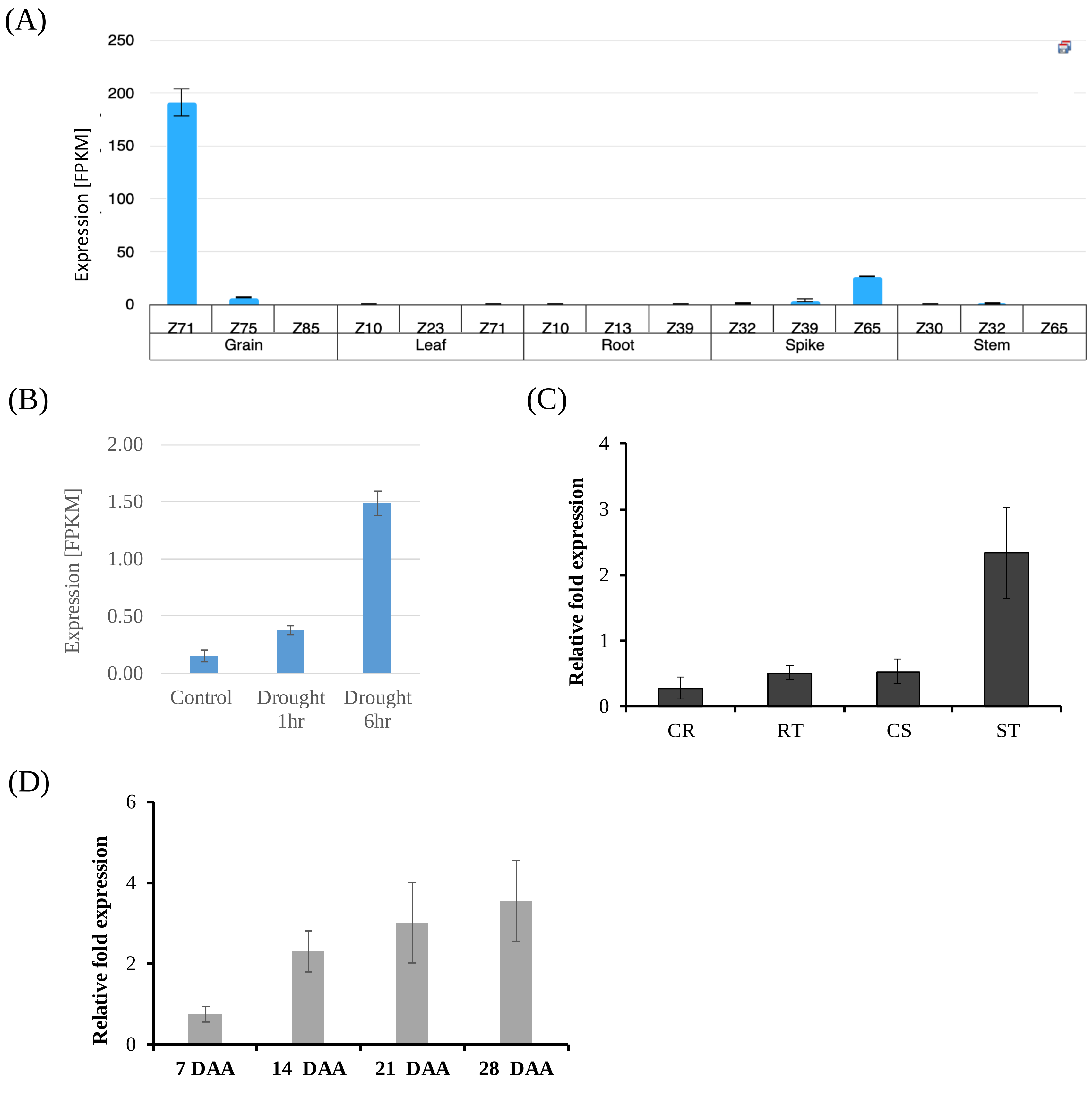


**Supplementary Figure S7: Expression of TaFLA7 in wheat.** (A) Relative gene expression level during plant development (at Zadosky scale; Exvip database; https://dubcovskylab.ucdavis.edu/expression_data/). (B) Expression of the *TaFLA7* after 1 and 6 days of drought stress. (C) The expression profiles of the FLA7 in roots and shoots tissue of wheat seedlings collected at 6 days under drought stress n=3 biological replicates. (D) Relative fold expression analysis of *TaFLA7* during grain development (days after anthesis-DAA).

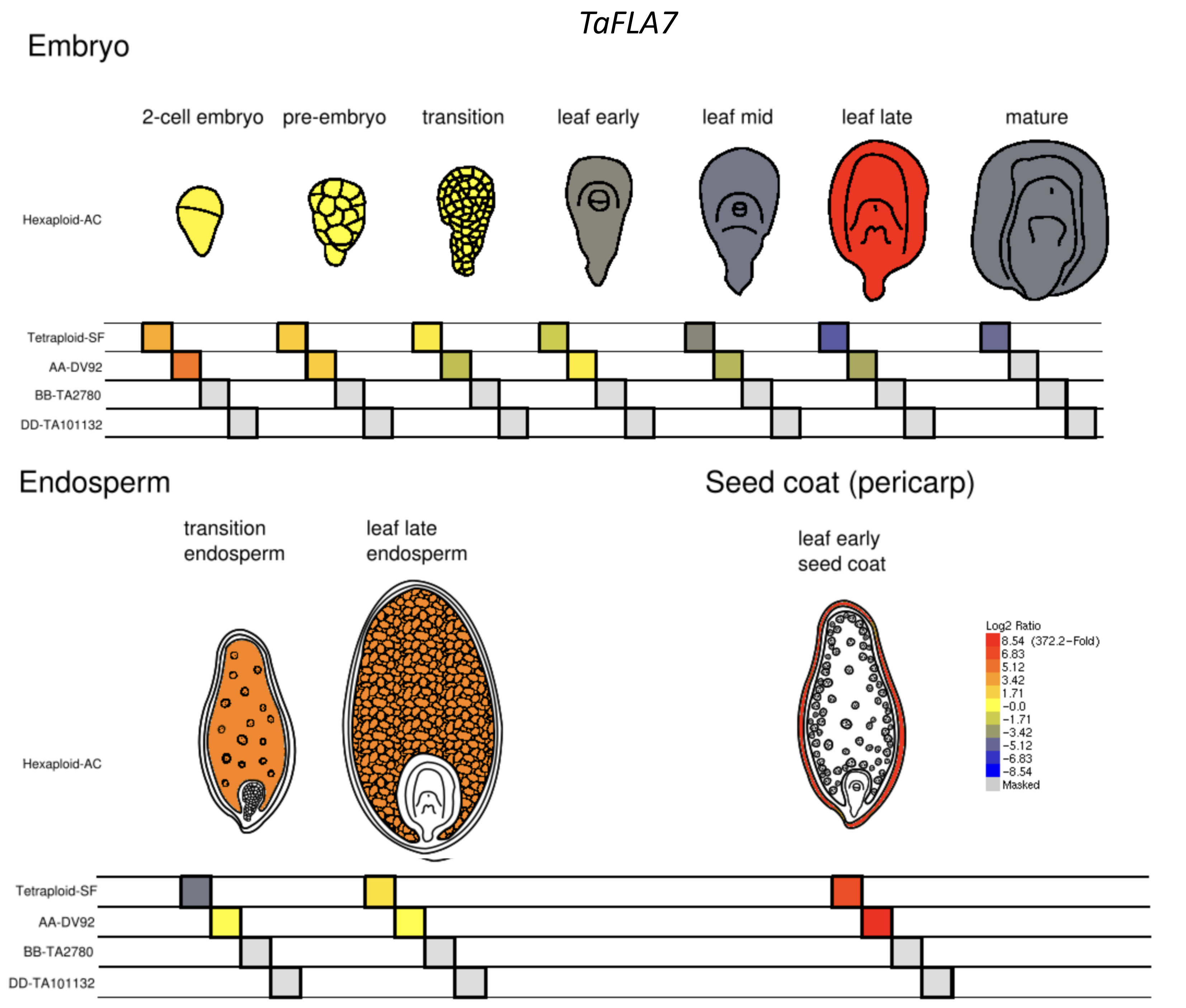


The authors used a Illumina Hi-seq sequencing platform and selected seven stages of embryo starting from fertilization to mature embryo, the selected key embryo development stages were determined by overall morphologies of the developing grass grains in relation to its main developmental phases with some modifications. These key stages including: two cell (E1), pre-embryo (E2), transition (E3), leaf early(E4), leaf middle(E5), leaf late(E6) and mature embryo(E7); two stages of isolated endosperm including: transition stage endosperm (E8) and leaf late stage endosperm (E9); and one stage of seed coat: leaf early stage pericarp) (E10) from diploid T. monococcum (DV92-DV), Ae. Speltoides (TA2780-SP) and Ae. Tauschii(TA101132-TA), tetraploid (T. durum, Canadian Cultivar, Strong Field-SF) and hexaploid (T. aestivum , Canadian Cultivar, AC Barrie-AC) for RNA-Seq analysis.

**Supplementary Figure S8:** Expression analysis of *TaFLA7* in different wheat grain tissue. eFP wheat browser (https://bar.utoronto.ca/efp_wheat/cgi-bin/efpWeb.cgi) was used to plot the data (using RNAseq data from *Triticum aestivum* cv. Azhurnaya spring wheat ) the heat map indicates the change in the gene expression level as $Log_2$ ratio.

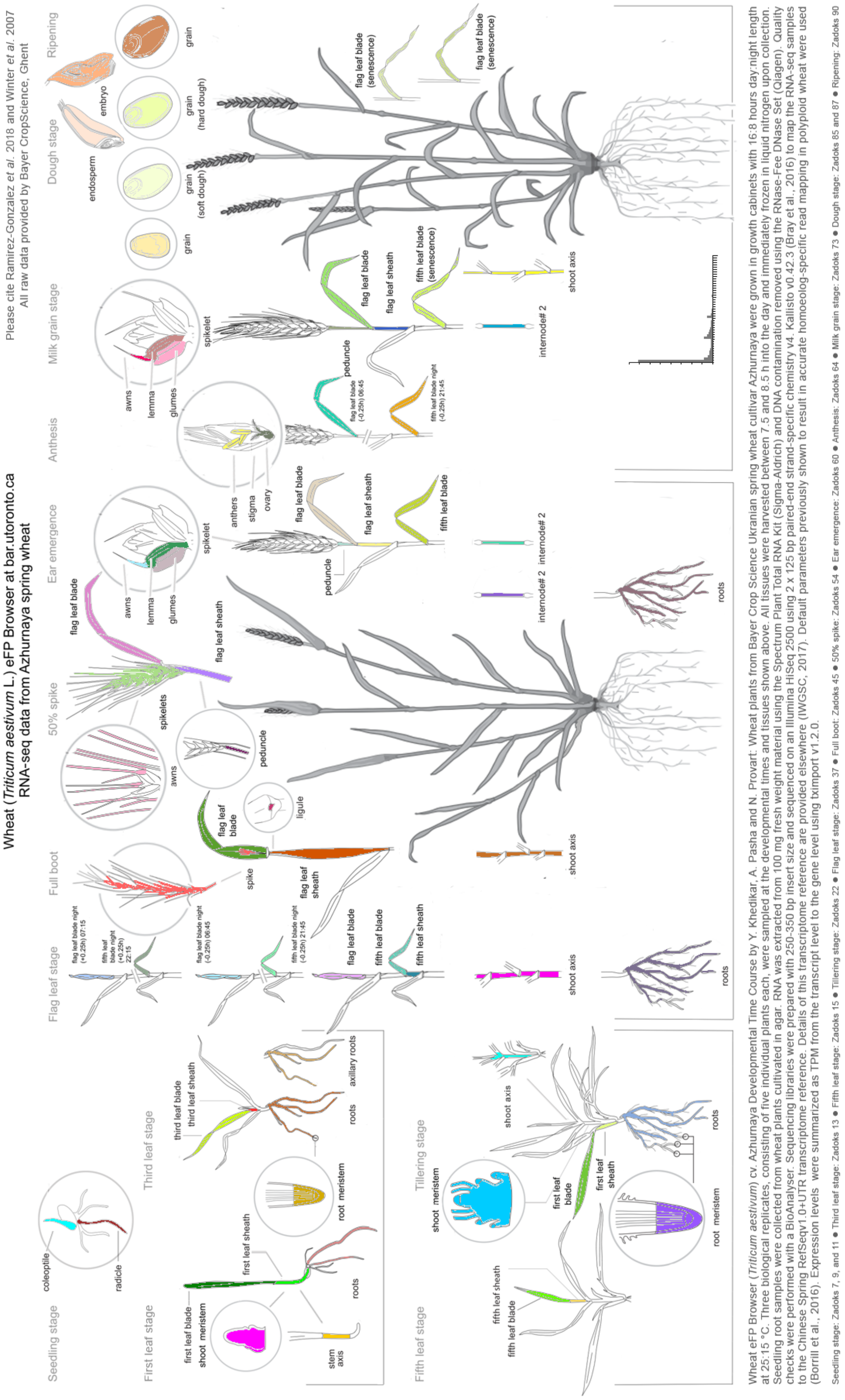

**Supplementary Figure S9:** Expression analysis of *TaFLA7* during wheat developmental stages and in various tissue. eFP wheat browser (https://bar.utoronto.ca/efp_wheat/cgi-bin/efpWeb.cgi) was used to plot the data (using RNAseq data from *Triticum aestivum* cv. Azhurnaya spring wheat ) the heat map indicates the change in the gene expression level as $Log_2$ ratio.
